# Population-specific patterns of toxin sequestration in monarch butterflies from around the world

**DOI:** 10.1101/2021.10.15.464593

**Authors:** Micah G. Freedman, Sue-Ling Choquette, Santiago R. Ramírez, Sharon Y. Strauss, Mark D. Hunter, Rachel L. Vannette

## Abstract

Animals frequently defend themselves against predators using diet-derived toxins. Monarch butterflies are a preeminent example of toxin sequestration, gaining protection via cardenolides in their milkweed hosts. Few studies have considered genetic variation in sequestration ability, in monarchs or other species. Here, we use two approaches to study natural selection on cardenolide sequestration in monarchs. First, we conducted a reciprocal rearing experiment with six monarch populations and six associated host species from around the world to determine whether sequestration is higher in monarchs reared on sympatric host species. Second, we compared sequestered cardenolides in wild-caught monarchs from Guam—an island where bird predators have been functionally extirpated for >40 years—to a nearby island with intact birds. We found substantial genetic variation in sequestration ability, though no consistent sequestration advantage in sympatric combinations. One monarch population from Puerto Rico showed greatly reduced sequestration from *Asclepias syriaca*, likely reflecting a lack of evolutionary association with this host. Monarchs from Guam showed reduced sequestration from *A. curassavica*, both in a cross-island comparison and when reared under controlled conditions. Our results suggest that processes involved in toxin sequestration are subject to natural selection and may evolve in response to contemporary changes in species interactions.

## Introduction

The use of diet-derived toxins as a defense against higher trophic levels is common across the tree of life (Brodie 2009) and has been documented in taxa as diverse as snakes (Hutchinson et al. 2007), poison dart frogs (Santos et al. 2003), and African crested rats (Kingdon et al. 2012). Toxic prey often gain protection against predation through the process of sequestration, defined as the selective uptake, transport, modification, storage, and deployment of secondary compounds (Heckel 2014). Variation in toxin sequestration behavior is often studied across species in phylogenetic comparative contexts (e.g., Engler-Chaouat and Gilbert 2007; Petschenka and Agrawal 2015) or within species in relation to prey availability (e.g., McGugan et al. 2016; Yoshida et al. 2020). However, we still have a limited understanding for how contemporary species interactions—both top-down (predators) and bottom-up (prey)—may exert natural selection on toxin sequestration behavior.

Herbivorous insects provide many of the clearest instances of toxin sequestration behavior (Petschenka and Agrawal 2016; Beran and Petschenka 2022), including sequestration of iridoid glycosides by nymphalid butterflies (Bowers and Puttick 1986), glucosinolates by flea beetles (Beran et al. 2014), pyrrolizidine alkaloids by arctiid moths (Von Nickisch-Rosenegk and Wink 1993), and aristolochic acids by swallowtail butterflies (Fordyce and Nice 2008). Numerous studies have documented variation in sequestration of defensive compounds from populations across the geographical range of species (e.g., Brower and Moffitt 1974; Gardner and Stermitz 1988), although these differences are usually attributed to differences in host plant availability. By contrast, relatively little research has focused on intraspecific genetic variation in the propensity to sequester dietary toxins (but see Müller et al. 2003; Fordyce and Nice 2008).

Monarch butterflies (*Danaus plexippus*) are perhaps the single best-studied example of a toxin-sequestering animal. Monarch larvae feed on milkweeds (Apocynaceae: Asclepiadoideae) and incorporate toxic cardiac glycosides (cardenolides) from these hosts that remain in their tissue throughout development (Brower et al. 1967, Reichstein et al. 1968, Roeske et al. 1976, Agrawal et al. 2017). Cardenolides sequestered by monarchs confer protection against bird predators, as demonstrated in the iconic series of experiments by Lincoln Brower and colleagues (Brower et al. 1968; Brower et al. 1972; Brower and Moffitt 1974) and the associated images of a vomiting blue jay. Sequestered cardenolides may also deter invertebrate predators (Rayor et al. 2004) and parasitoids (Stenoien et al. 2019).

Despite research into variation in sequestration across monarch tissues (Brower and Glazier 1975; Frick and Wink 1995), across their ontogeny (Jones et al. 2019), throughout their migratory cycle (Malcolm and Brower 1989), and across the broader phylogeny of milkweed butterflies (Petschenka et al. 2013, Petschenka and Agrawal 2015, Karageorgi et al. 2019), little is currently known about how natural selection may shape sequestration abilities over contemporary time scales. One approach that could improve our understanding of the selective forces operating on sequestration involves using geographically disparate populations of monarchs with divergent host plant assemblages to test for enhanced sequestration ability in butterflies reared on sympatric host species (Figure 1A). Monarch populations around the world show some evidence for local adaptation to their available host plants based on larval growth rate (Freedman et al. 2020a), as well as subtle variation in the terminal domain sequences of cardenolides’ target enzyme (the sodium-potassium pump, Na^+^/K^+^-ATPase) (Pierce et al. 2016). Here, we predict that monarch populations should have a sequestration advantage when reared on sympatric host species, and that patterns of cardenolide sequestration should reflect the history of association between each monarch-milkweed pair.

**Figure 1.**
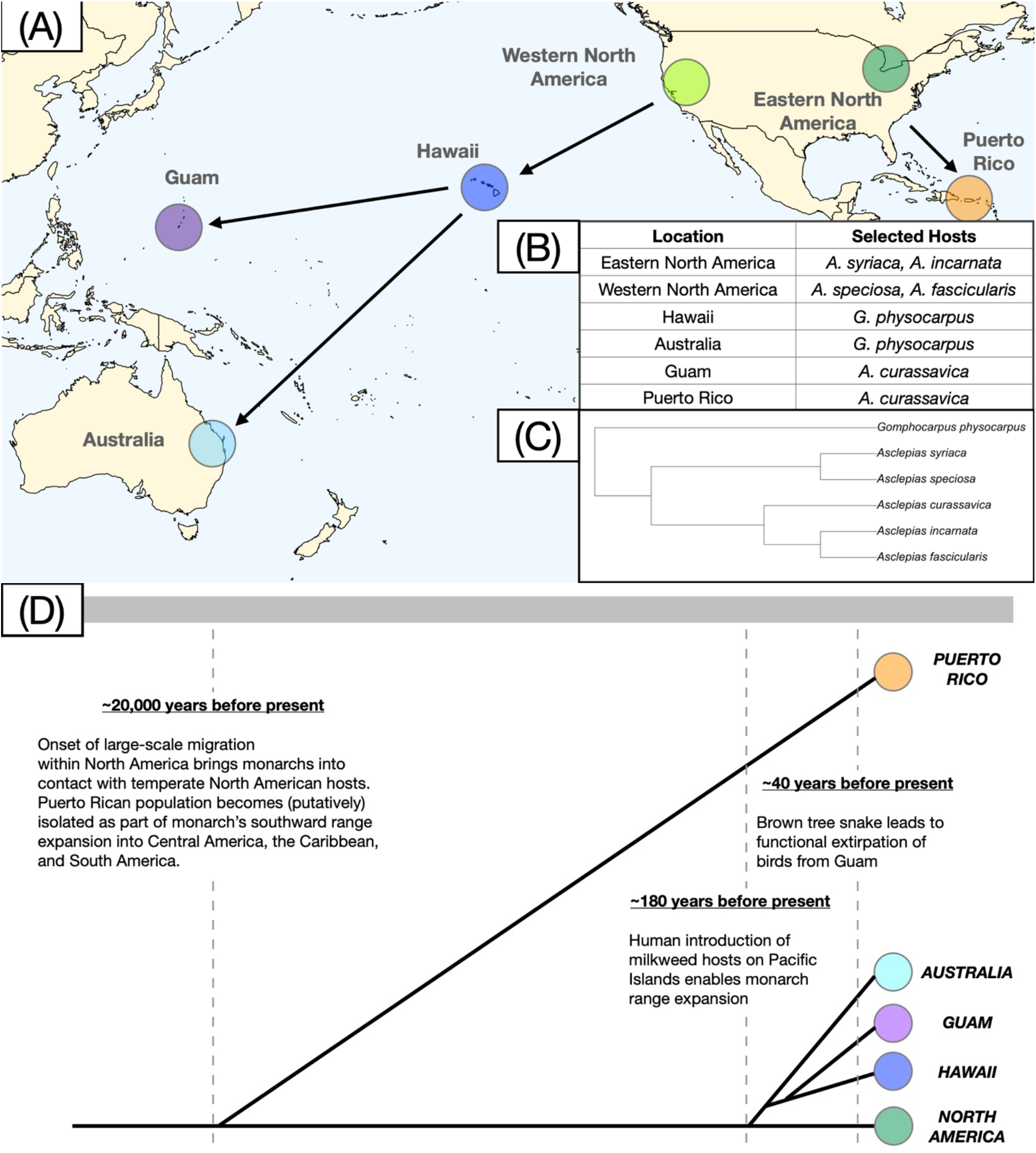
(**A**) Monarch populations and their associated host plants. Arrows indicate the direction of expansion out of the ancestral North American range. Pacific Island populations are part of a single westward expansion event, while Puerto Rican monarchs are part of an independent southward expansion event. Selected hosts for each population are considered sympatric in all analyses. (**B**) Summary of monarch populations and their sympatric hosts used in current study. (**C**) Phylogeny of milkweed species used in current study, recreated from Agrawal and Fishbein (2008). (**D**) Phylogram showing relatedness among populations and approximate timing of major events discussed throughout the manuscript. Note that timeline is not calibrated and that Eastern and Western North American samples are shown as part of a single North American population but are treated as separate populations with distinct host plant assemblages.

Another approach to studying selection on sequestration behavior involves using natural variation in exposure to predation. Although the protective benefits of sequestered cardenolides as a defense against predation have been shown (e.g., Fink and Brower 1981, Rayor et al. 2004), there are potential costs: sequestered cardenolides can inhibit even the highly insensitive Na^+^/K^+^-ATPase of monarchs (Petschenka et al. 2018; Züst et al. 2019; Agrawal et al. 2021) and impose physiological costs associated with flight metabolism (Pocius et al. 2022) and oxidative stress (Blount et al. 2021). These costs may be particularly acute on hosts such as *A. curassavica* (Agrawal et al. 2021, Pocius et al. 2022), from which monarchs sequester high concentrations of especially potent cardenolides. Cardenolide sequestration may therefore involve balancing anti-predator benefits with physiological costs. Accordingly, we predict that monarchs with reduced exposure to predators should sequester lower concentrations of cardenolides.

In this study, we conduct a fully reciprocal rearing experiment using six monarch populations and six associated host plant species from around the world and measure cardenolide sequestration in a set of 440 butterflies. We test for local adaptation in sequestration ability by determining whether monarchs have a sequestration advantage when reared on their sympatric host species. We also assess tradeoffs in sequestration ability across hosts, as well as inherent variation in sequestration among monarch populations and host plants. Finally, we focus on one monarch population from Guam that has lost its bird predators over recent evolutionary history to determine whether this loss of predators is associated with a reduction in cardenolide sequestration.

## Methods

### Study system and natural history

Monarch butterflies are best-known from their ancestral range in North America, where they migrate seasonally and feed on more than 40 milkweed host species (Malcolm and Brower 1986; Xerces Society 2018). Over recent evolutionary history, monarchs have greatly expanded their geographic range and are now established in locations throughout Central and South America, the Caribbean, the Pacific, and the Atlantic (Vane-Wright et al. 1993, Pierce et al. 2014; Zhan et al. 2014), with Pacific and Atlantic populations likely becoming established in the last ~180 years (Zalucki and Clarke 2004; Freedman et al. 2020b). Nearly all recently established monarch populations are non-migratory and breed year-round on restricted assemblages of host plants (Pierce et al. 2016; Freedman et al. 2020a). Monarchs have little coevolutionary history with many of the host plants in their introduced range, and host plant species available to monarchs in locations throughout the Pacific and Atlantic—primarily *A. curassavica*, but also *Gomphocarpus spp*. and *Calotropis spp*.—are themselves recent introductions from subtropical Africa, India, and the Neotropics.

Monarch butterflies are subject to predation throughout their lifetime: major predators of larvae and eggs include Tachinid flies (Oberhauser et al. 2017), Polistine wasps (Baker and Potter 2020), ants (Calvert 2004), and various opportunistic generalists including earwigs (Hermann et al. 2019). Adults are thought to be primarily attacked by birds (Calvert et al. 1979; Brower 1988; Groen and Whiteman 2021), although mice are likely also a major source of adult predation, especially at overwintering locations (Glendinning and Brower 1990; Weinstein and Dearing 2022).

One monarch population that we sampled (Guam) is unique in that butterflies there have been functionally released from bird predation since the late 1980s. The loss of insectivorous birds from Guam resulted from the introduction of the brown tree snake (Savidge 1987) and is associated with changes in the island’s trophic ecology (Rogers et al. 2012). Thus, by comparing sequestration in monarchs from Guam to locations with more intact assemblages of bird predators, it is possible to test for an association between exposure to bird predation and sequestration, though with the major caveat that the population-level bird exclusion treatment in Guam is not replicable.

### Experiment 1: Reciprocal Rearing

Over the course of two years, we conducted a fully factorial rearing experiment using six populations of monarchs from around the world and their associated host plants (Figure 1A-C). This experiment is the same as the one described in Freedman et al. (2020a), although here we focus on cardenolide sequestration as the phenotype of primary interest rather than larval growth rate. We used the following six host plant species: *A. curassavica*, *A. incarnata*, *A. fascicularis*, *A. speciosa*, *A. syriaca*, and *G. physocarpus* (Figure 1C). Host plants were grown from seed in 1-gallon pots in two greenhouses; for details of host plant provenance, see Table S1. Monarchs were collected in the field from six global sites as gravid adult females and returned live to UC Davis in glassine envelopes, where females laid eggs on cut stems of *A. curassavica*. Within 12 hours of hatching, we transferred neonate larvae onto a randomly assigned host plant using a paintbrush, typically adding 5 larvae per plant. When possible, we used a balanced design that assigned larvae from a single maternal family to all possible host plants (Table S2). We then used mesh sleeves to restrict larvae to a single live host plant. For each caterpillar, we recorded its mass after 8 days, the number of days until pupation, and the number of days until eclosion; all of these data are reported in Freedman et al. (2020a). Because multiple butterflies were reared from individual host plants, we were unable to keep track of individual identity and match butterflies to their larval mass on day Full details of plant propagation and caterpillar rearing are reported in Supplementary Appendix 1.

#### Tissue collection and processing

We extracted cardenolides from milkweed leaf discs and entire butterfly hindwings. Adult monarchs store cardenolides primarily in their wings and integument, with wing cardenolides thought to be an adaptation for deterring bird predation (Brower and Glazier 1975). We chose to measure cardenolides from monarch wings for a number of reasons: previous work has shown that wings are the tissue with the highest concentrations of cardenolides (Brower and Glazier 1975, Fink and Brower 1981), wing cardenolide concentrations are highly correlated with cardenolide concentrations in other tissues (Fink and Brower 1981, Figure S1), and wings are the tissue most likely to be used by predators to assess monarch palatability (Fink and Brower 1981). For a full description of cardenolide extraction methods, see Supplementary Appendix 2. In total, we collected data from 183 leaf samples and 451 wing samples. All of our analyses use total cardenolide concentrations in adult hindwings as our response variable.

#### Cardenolide identification and quantification

We used ultra-performance liquid chromatography (UPLC) to separate cardenolides based on their polarity; compounds with early retention times are more polar than those that elute later (see Figure 2A for chromatograms). Peaks with absorbance spectra between 216-222 nm were considered to be cardenolides (Zehnder and Hunter 2007; Jones et al. 2019). Across all samples, we identified 70 distinct peaks, each of which should correspond to a unique cardenolide compound. We note that some of these peaks are likely constituent fragments of larger, more intact cardenolides; for example, the widespread cardenolides calotropin, calotoxin, calactin, and uscharin share a common aglycone precursor (calotropogenin). In order to verify the identity of some major sequestered compounds, we tested authentic standards for the compounds calactin, calotropin, and frugoside—reported to be the three major compounds sequestered from *A. curassavica* (Agrawal et al. 2021)—as well as aspecioside, reported to be a major sequestered compound from *A. syriaca* (Seiber et al. 1986; Malcolm et al. 1989; Agrawal et al. 2022) (Table S3). Authentic standards were provided by A. Agrawal and are the same as those used in Agrawal et al. (2021). Cardenolide peak areas were measured at 218 nm and integrated using Chromeleon™ software (Thermo-Fisher). Each sample was prepared with a digitoxin internal standard (Sigma-Aldrich) added at a known concentration to allow for quantification.

**Figure 2.**
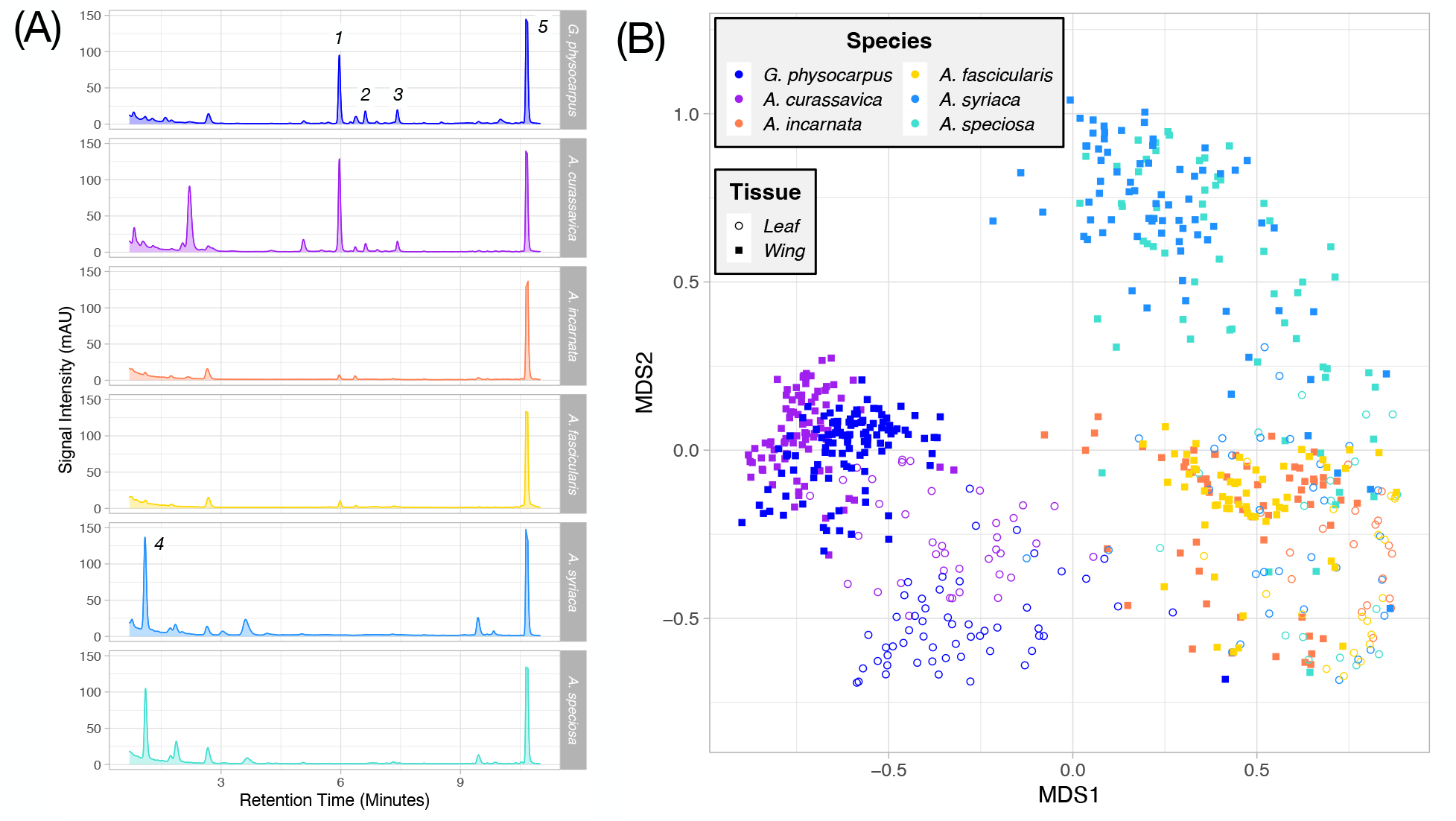
**(A)** Example of chromatograms showing sequestered cardenolides in monarch hindwings. Each panel reflects a butterfly from one of the six milkweed species used during rearing. Retention times correspond to compound polarity, with more polar compound eluting first and less polar compounds eluting last. Numbered peaks were verified with authentic standards and are as follows: 1 = frugoside, 2 = calotropin, 3 = calactin, 4 = aspecioside, 5 = digitoxin (internal standard). Note that the y-axis is truncated and does not show the true values for the internal standard (digitoxin – 0.15 mg/mL), which elutes around 10.8 minutes and was generally the largest peak in each sample. **(B)** NMDS plot of leaf and wing tissue. Note the similarity in the profiles of sequestered compounds from *A. curassavica* and *G. physocarpus*, as well as *A. speciosa* and *A. syriaca*.

### Data Analysis: Experiment 1

#### Multivariate disparity in cardenolide profiles

We plotted raw data to explore variation in patterns of sequestration across host plants and monarch species. We visualized multivariate disparity in cardenolide profiles of wings and leaf tissue using non-metric multidimensional scaling implemented in the R package ‘vegan’ (v2.5-7) (Oksanen et al. 2020). We used PERMANOVA (implemented using the adonis2 function, a matrix of Bray-Curtis dissimilarities, and with 1000 permutations) within each milkweed species to test whether leaf and corresponding wing samples had significantly different cardenolide profiles. We also analyzed multivariate disparity in sequestered cardenolides using PERMANOVA and a model that considered milkweed species, monarch population, and their interaction as predictors.

#### Measuring GxE interactions and local adaptation for sequestration

To test for quantitative variation in cardenolide sequestration across host species and monarch populations, we used linear mixed models implemented in the lme4 package (Bates et al. 2015) in R version 4.0.3 (R Development Team). Since sequestration amounts were consistently low across all populations for two species (*A. fascicularis* and *A. incarnata*) (see Figure 3) (also see Malcolm 1994), we restricted these analyses to only monarchs reared on the remaining four milkweed species (n = 327). First, to quantify GxE interactions and measure variation across each *monarch population x milkweed species* combination, we fit a model of the form:

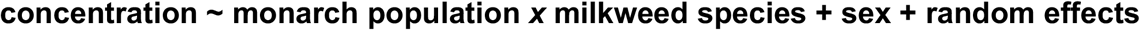

where concentration refers to the hindwing cardenolide concentration of an individual butterfly. We included a categorical factor for butterfly sex to account for potential differences males and females in sequestration (Brower and Glazier 1975). We included random effects for monarch maternal family of origin as well plant ID nested within plant population of origin. Model results were summarized using Type III ANOVAs implemented in the ‘car’ package (Fox and Weisberg 2019). We assessed post-hoc pairwise differences between monarch populations and milkweed species using TukeyHSD tests implemented in the ‘multcomp package’ (Hothorn et al. 2008).

**Figure 3.**
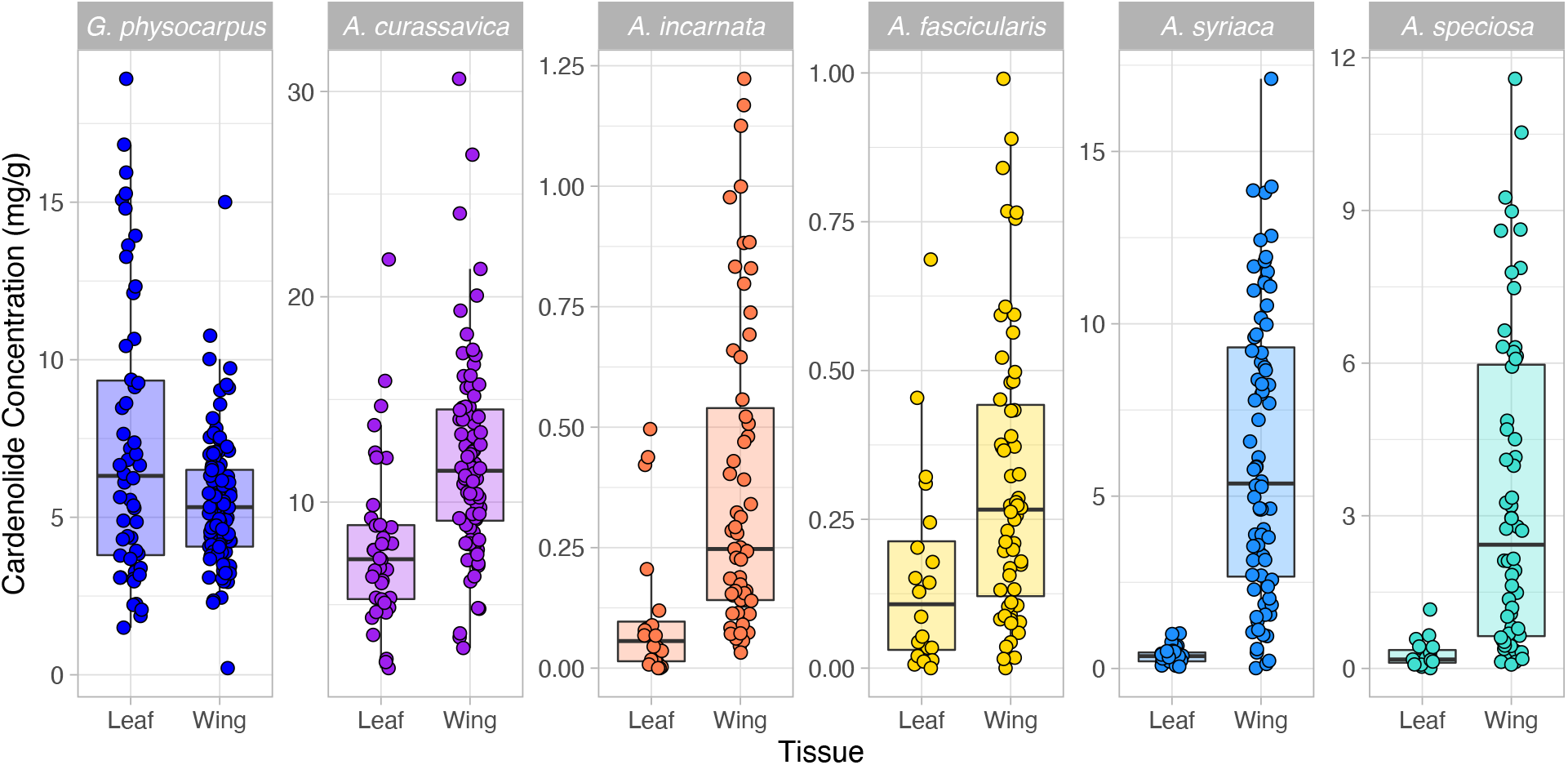
Boxplots showing cardenolide concentrations (expressed as milligrams of cardenolide per gram of oven-dried leaf or wing tissue) for leaf and wing tissue of each milkweed species. Here, each point corresponds to either a single plant tissue sample or a single butterfly hindwing. Note that y-axes differ substantially between species. Figure S3 shows the association between individual butterflies and their natal host plants.

To test for local adaptation in cardenolide sequestration, we fit a model of the form:

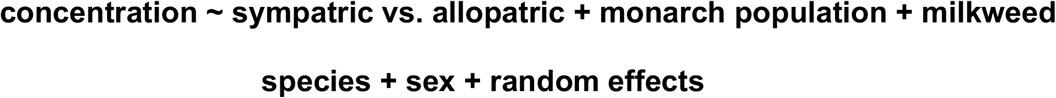

The formulation of this model is the same as the previous model, but instead of including interaction terms for each *monarch population x milkweed species* combination, it features a term that describes whether this combination is sympatric or allopatric. In this model, the primary effect of interest is sympatry vs. allopatry: a significant positive intercept for sympatric combinations is diagnostic of local adaptation (Blanquart et al. 2013).

To account for the possibility that an individual butterfly’s level of sequestered cardenolide varied as a function of the leaf cardenolide content of its specific host plant (rather than just host species identity), we also tested a model that included plant cardenolide concentration as a predictor variable. We then compared this model to an equivalent model without plant cardenolide concentration using ΔAIC scores.

### Experiment 2: Comparing Sequestered Cardenolides in Guam and Rota

In July 2015, we collected wild monarchs from Guam (n = 54), Rota (n = 27), and Saipan (n = 2) in the Mariana Islands (Figure 5A). Birds have been extirpated from Guam since the 1980s, while Rota and Saipan both have mostly intact insectivorous bird assemblages. Monarchs from Guam and Rota were collected from host plant patches consisting entirely of *A. curassavica*, and we also collected a small number of *A. curassavica* samples from each island. As with greenhouse-reared butterflies, we used a single hindwing from each butterfly to measure sequestered cardenolides. Cardenolide quantification was performed similarly to the methods described above but using a separate HPLC instrument at the University of Michigan.

After quantification, we inspected multivariate cardenolide profiles for butterflies from each island. Butterflies from Saipan were excluded from subsequent analysis due to their small sample size and disparate cardenolide profiles (Figure S2). For butterflies from Guam and Rota, we created an index of wing wear and assigned each individual a value from 1-5 (Table S4) (see Malcolm et al. 2018). This was wing wear value was used as a proxy for butterfly age, which is negatively correlated with cardenolide content (Tuskes and Brower 1978). We compared total cardenolide concentrations (natural log transformed to account for the non-linear decrease in cardenolides across wing wear values) in butterflies from Guam and Rota using a linear model of the form:

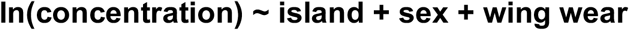

Our approach to comparing monarchs from Guam and Rota has a number of important caveats. First, it necessarily involves a functional sample size of n = 1, as Guam is the only bird-free island from which we could sample. This is an inherent weakness of this system that cannot be avoided. Second, we attempted to collect live butterflies from Rota to include in controlled rearing experiments in 2018 but were unable to locate any. Thus, we can only compare wild-caught butterflies from each island, which does not allow for us to formally disentangle genetic from environmental sources of variation in sequestered cardenolides. However, we feel fairly confident attributing any observed differences in wing cardenolide concentrations to genetic differences between islands for the following reasons: (1) all monarchs from Guam and Rota appear to have developed on the same host, *A. curassavica* (Figure S2); (2) in our greenhouse rearing experiment, plant cardenolide concentrations in *A. curassavica* did not positively correlate with butterfly wing concentrations (Figure S7), suggesting that any observed differences in wild-caught butterflies are unlikely to be driven by variation in host plants; (3) despite their proximity, monarchs from Guam and Rota show strong genome-wide patterns of differentiation (Hemstrom et al. 2022), highlighting the potential for phenotypic divergence.

## Results

### Overall patterns of variation in milkweed and monarch cardenolides

Milkweed species varied greatly in their cardenolide composition (Figure 2A, 2B) as well as their average cardenolide concentration, ranging from as low as 0.11 ± 0.03 mg/g (*A. incarnata*) to as high as 7.86 ± 0.66 mg/g (*A. curassavica*) (Figure 3). Monarchs, regardless of population of origin, had the highest levels of sequestered cardenolides on *A. curassavica* (12.11 ± 0.53 mg/g) and the lowest on *A. fascicularis* (0.31 ± 0.03 mg/g) (Figure 3). Monarchs reared on *A. syriaca* had hindwing cardenolide concentrations that were more than 12 times higher than their natal host plant tissue; by contrast, for monarchs reared on *G. physocarpus*, hindwing concentrations were 1.3 times lower than corresponding leaf tissue (Figure 3). Across all species and populations, female monarchs sequestered slightly more than males, although this difference was not significant (t = 1.688, p = 0.091) (Table S9). The polarity index of sequestered cardenolides varied strongly across species: in general, monarchs reared on *A. syriaca* and *A. speciosa* sequestered primarily polar cardenolides, while the subset of sequestered cardenolides on other species was predominantly compounds with intermediate polarity (Figure 2A, Figure S4).

Across all milkweed species, the composition of cardenolides present in leaves was significantly different from the composition of sequestered cardenolides (Figure 2B; Table S5). Calactin, calotropin, and frugoside were present in monarchs reared on *A. curassavica* and *G. physocarpus*, and together comprised approximately 50% of the total amount sequestered in hindwings for both species (Table S6). Aspecioside was the predominant compound sequestered from both *A. syriaca* and *A. speciosa* (Table S6). Within milkweed species, concentrations of individual sequestered cardenolides were generally positively correlated (Figure S5). The overall composition of sequestered cardenolides was most strongly determined by milkweed species identity (F = 119.49, R^2^ = 0.494), followed by monarch population (F = 4.77, R^2^ = 0.033), and finally the interaction between them (F = 2.85, R^2^ = 0.059) (Figure 2B; Figure S6; Table S7).

Within most milkweed species, there was not a strong correspondence between leaf cardenolide concentrations and wing cardenolide concentrations: only one milkweed species (*A. speciosa*) showed a significant positive correlation between leaf and wing concentrations (Figure S7), and a model that did not include plant-level leaf chemistry was favored over one that did (Table S8). Thus, all reported analyses are based on untransformed monarch hindwing cardenolide concentrations, irrespective of leaf cardenolide concentrations from the corresponding natal host plant.

### GxE interactions for sequestration

We found strong support for GxE interactions in sequestration ability, with monarch populations varying substantially in their ability to sequester across milkweed species (X^2^ = 77.6, d.f. = 15, p < 0.001) (Figure 4A; Table S9). This pattern was driven most strongly by cross-host sequestration differences in monarchs from Puerto Rico. Puerto Rican monarchs sequestered 1.37 times more from *A. curassavica* and 1.46 times more from *G. physocarpus* than other populations, yet 4.96 times less from *A. speciosa* and 5.83 times less from *A. syriaca* (Figure 4A, 4C; Figure S8). The polarity index of cardenolides sequestered by Puerto Rican monarchs on *A. syriaca* and *A. speciosa* was significantly lower than for all other populations (t = −6.86, p < 0.001; Figure S4), and the sequestration profile of Puerto Rican monarchs was distinct from other monarch populations on *A. syriaca* (Figure S6).

**Figure 4.**
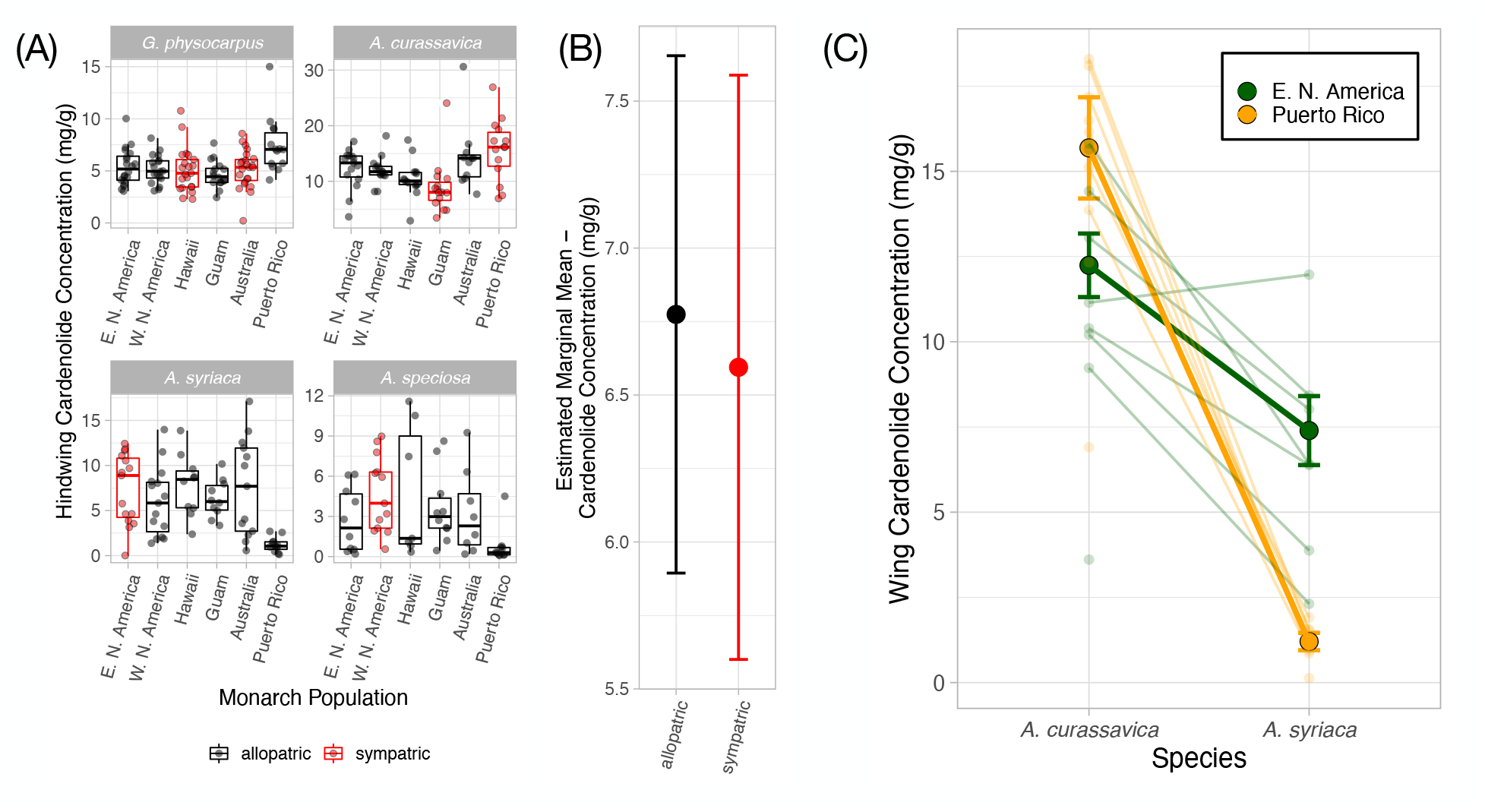
**(A)** Boxplots showing levels of sequestered cardenolides across each *monarch population x host species* combination. Each point corresponds to a single butterfly. Sympatric combinations are shown in red and allopatric combinations in black. **(B)** Model-averaged estimates of overall cardenolide sequestration across all sympatric and allopatric combinations. Monarchs reared on sympatric hosts had no sequestration advantage. **(C)** Reaction norm plot showing GxE interactions for sequestration in Puerto Rican and North American monarchs reared on *A. curassavica* and *A. syriaca*. Dark points correspond to population-level means for each combination; lighter points and faint lines show means and reaction norms for each individual maternal family within each population. For this comparison, the *monarch population x host species* interaction term is highly significant (t = 3.931, p < 0.001).

**Figure 5.**
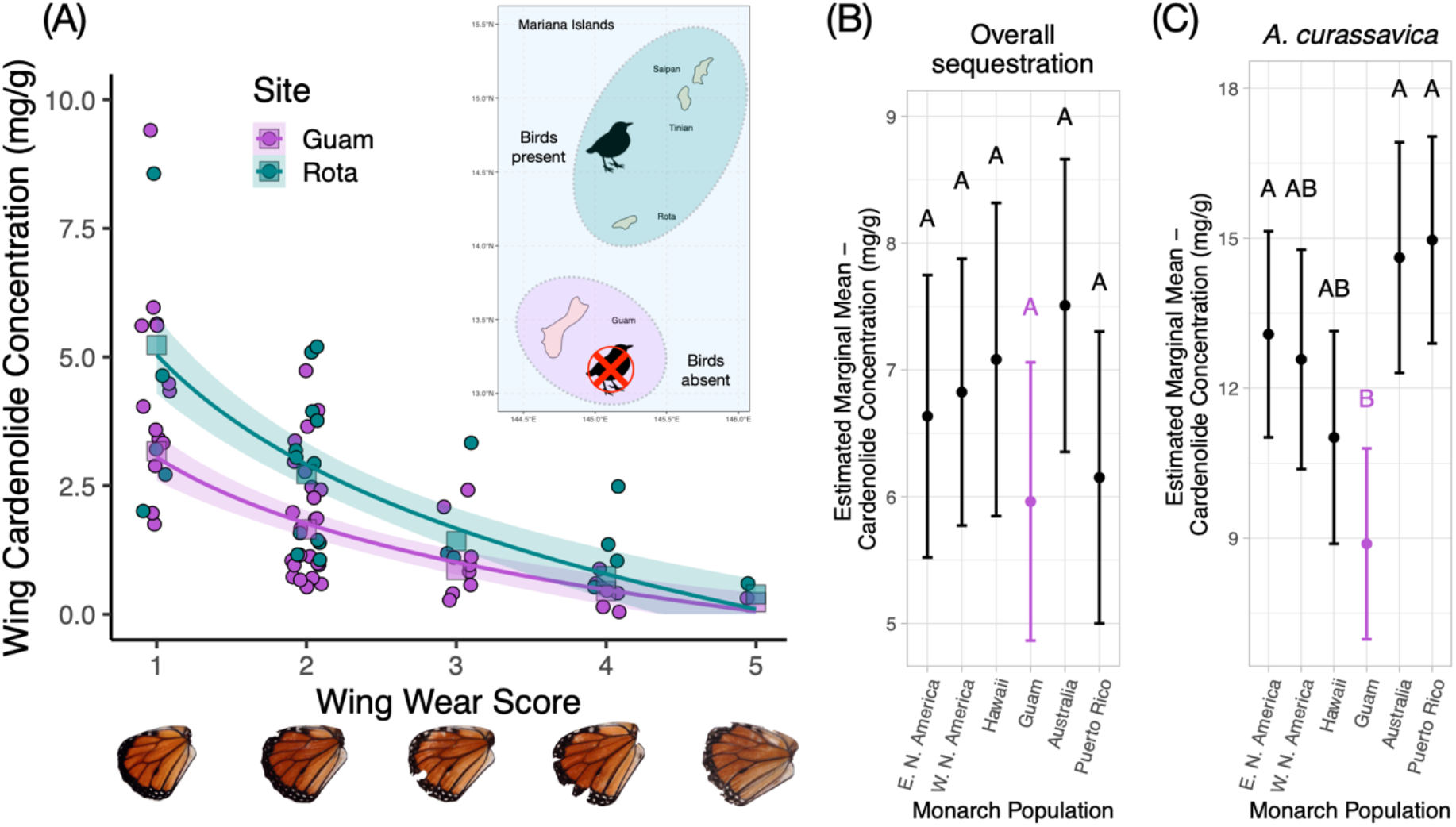
**(A)** Comparison of hindwing cardenolides in wild-caught monarchs from Guam and Rota across the range of wing wear scores. Examples of wings assigned to each wing wear category are shown along the X axis. Curves depict model-fitted estimates (squares) and 95% confidence intervals. Inset map shows the Mariana Islands. (**B**) Overall levels of cardenolide sequestration across all host species for each monarch population tested in the greenhouse rearing experiment. Monarchs from Guam had the lowest population-specific intercept, but this value was not significantly different from other populations after correcting for multiple comparisons. (**C**) Levels of cardenolide sequestration on *A. curassavica* in the greenhouse rearing experiment. Here, monarchs from Guam sequestered significantly less than monarchs from Eastern North America, Australia, and Puerto Rico.

### Local adaptation for sequestration

Despite the strong GxE pattern of sequestration in our data, there was no support for local adaptation in sequestration ability (X^2^ = 0.16, d.f. = 1, p = 0.687), with roughly equivalent levels of sequestration in sympatric and allopatric *population x host* combinations (Figure 4A, 4B; Table S10). Accounting for development time did not meaningfully impact any of our inferences (Figure S9), and we did not find a strong correlation between development time and the concentration of hindwing cardenolides (Figure S10). Maternal families within populations varied substantially in their propensity to sequester cardenolides (Figure S11).

### Sequestration in monarchs from Guam

Wild-caught monarchs from the bird predation-free island of Guam has significantly lower concentrations of hindwing cardenolides than monarchs from Rota (t = −3.119, p = 0.003) (Figure 5A). The degree of wing wear (a proxy for butterfly age) was by far the strongest predictor of butterfly hindwing concentrations (t = −9.164, p < 0.001) (Figure 5A), with older butterflies having lower cardenolide concentrations. After accounting for differences in wing wear and butterfly sex, Rota monarchs had hindwing cardenolide concentrations (2.30 ± 1.14 mg/g) that were 1.65 times higher than monarchs from Guam (1.39 ± 1.09 mg/g).

Among the six monarch populations reared under controlled conditions in the greenhouse, Guam had the lowest population-specific intercept for overall cardenolide sequestration (Figure 5B). Notably, the pattern of reduced sequestration by monarchs from Guam was most pronounced on *A. curassavica*, their sympatric host: Guam monarchs sequestered, on average, 33.1% fewer cardenolides on *A. curassavica* than other populations, and significantly less than populations from Australia, Eastern North America, and Puerto Rico on this host (Figure 5C; Table S11). However, this pattern of reduced sequestration was not detectable on other hosts, and the overall population-specific intercept for Guam was not significantly lower than for other populations (Figure 5B).

## Discussion

We found strong evidence for GxE interactions in sequestration ability, suggesting that monarchs and other taxa that sequester dietary toxins may show spatially structured genetic variation in their propensity to sequester. This GxE pattern in cardenolide sequestration was primarily driven by a single monarch population from Puerto Rico: these monarchs sequestered higher cardenolide concentrations than all other populations when reared on *A. curassavica* and *G. physocarpus*, but substantially lower concentrations when reared on *A. syriaca* and *A. speciosa* (Figure 4A; Table S12).

The most likely explanation for the Puerto Rican population’s reduced sequestration on *A. syriaca* and *A. speciosa* is a lack of evolutionary history with these hosts (Figure 1B). Divergence times between Puerto Rican monarchs and their migratory North American ancestors are uncertain but likely occurred within the last 20,000 years (Zhan et al. 2014), whereas other non-migratory populations included in this study were much more recently derived, likely diverging in the last 150-200 years (Zalucki and Clarke 2004; Freedman et al. 2020b). Thus, the lineage of Caribbean and South American monarchs that includes Puerto Rico may have a longer history of relaxed selection on sequestering from North American milkweeds, or alternatively may diverged prior to the onset of widespread adoption of *A. syriaca* and *A. speciosa* as hosts in North America. Recent evidence suggests that *A. syriaca* has undergone significant demographic expansions coinciding with post-glacial expansion (5-12 thousand years ago) and agricultural land use changes in North America (100-250 years ago) (Boyle et al. 2022). Under this scenario, Puerto Rican monarchs may have never evolved mechanisms to efficiently sequester a subset of distinctive cardenolide compounds (e.g., labriformin) present in widespread temperate North American milkweeds, including *A. syriaca*, *A. speciosa*, and *A. eriocarpa* (Nelson et al. 1981, Agrawal et al. 2022). Further research with additional monarch populations from the Caribbean and South America and/or additional North American milkweed species could help to resolve this question.

An alternative (but not mutually exclusive) explanation for the observed pattern of sequestration in Puerto Rican monarchs is a physiological tradeoff in sequestration ability, potentially driven by differences in the physical properties of cardenolides across milkweed host species. Puerto Rican monarchs sequestered high concentrations from *A. curassavica* and *G. physocarpus*, both of which are high cardenolide species and whose sequestration profiles are biased towards compounds with low to intermediate polarity (Roeske et al. 1976, Malcolm 1990). Interestingly, Puerto Rican monarchs sequestered higher cardenolide concentrations from *G. physocarpus* than any other population (Figure 4A; Table S12), despite little apparent history of association with this species, suggesting that feeding on *A. curassavica* or other high cardenolide hosts may have pre-adapted them to sequestering from the chemically similar *G. physocarpus*. By contrast, Puerto Rican monarchs sequestered very low concentrations of polar cardenolides from *A. syriaca* and *A. speciosa* that were readily sequestered by all other monarch populations (also see Seiber et al. 1986, Malcolm et al. 1989).

Despite finding evidence for unique sequestration behavior in monarchs from Puerto Rico, we did not find general evidence for a pattern of local adaptation in sequestration ability, with no overall support for greater sequestration from sympatric host plants across monarch populations. One possible reason for the lack of a sympatric sequestration advantage is that larval performance—including the process of sequestration—may be correlated across chemically similar host plants, even if they are geographically disparate and phylogenetically distant (e.g., Pearse and Hipp 2009). For example, the profile of cardenolides sequestered from *A. syriaca* and *A. speciosa* was nearly identical (Figure 2B; Table S6; Seiber et al. 1986), despite these two milkweed species having largely non-overlapping geographic ranges (Woodson 1954). Notably, we did not find evidence that monarchs from Hawaii or Australia have a sequestration advantage on *G. physocarpus*, despite apparently having >100 years of association with this host (Nelson 1993, Malcolm 1994). All derived Pacific Island populations (Hawaii, Australia, Guam) also retained their ability to sequester normally from ancestral North American hosts (*A. syriaca*, *A. speciosa*), even after spending as many as 1,500 generations isolated from these hosts.

Although monarchs did not show evidence of local adaptation in their sequestration behavior, this result may be biased by (1) relatively recent divergence times between most of the monarch populations that we tested (Puerto Rico being the exception); (2) strong dispersal capabilities in monarchs, which limits opportunities for specialization in allopatry; (3) the relatively limited diversity of cardenolides sequestered by monarchs. By contrast, sequestering species such as the strawberry poison-dart frog (*Oophaga pumilio*) that have limited dispersal capability, that show pronounced turnover in dietary composition over relatively small spatial scales, and that sequester more than 230 distinct alkaloid compounds from a range of functional classes might be stronger candidates for detecting local adaptation in sequestration ability (Saporito et al. 2007; Prates et al. 2019).

We found evidence for reduced cardenolide sequestration in monarchs from Guam, where birds have been functionally extirpated for the last ~40 years. In a comparison of wild-caught monarchs from Guam (birds absent) and nearby Rota (birds present), monarchs from Guam had significantly lower hindwing cardenolide concentrations. Guam monarchs also sequestered significantly less from their sympatric host plant (*A. curassavica*) than three other monarch populations, whereas sequestration was comparable across other hosts. This pattern is consistent with selection against the specific processes involved with sequestration from *A. curassavica* (e.g., the conversion of voruscharin into calotropin [Agrawal et al. 2021]), but not against other processes involved in the broader context of sequestration (e.g., multidrug transporter activity [Groen et al. 2017]). The observed reduction in sequestration from *A. curassavica*, but not other species, in monarchs from Guam accords with recent research showing that the physiological costs of sequestering from this species are especially pronounced (Agrawal et al. 2021; Blount et al. 2021; Pocius et al. 2022). Because it was only possible to test a single location with a long-term absence of bird predation, our ability to make broad inferences regarding predation intensity and selection for sequestration are limited. Still, the observation of reduced sequestration on a sympatric host, presumably under an altered predation regime, highlights the importance of considering higher trophic levels when forming predictions about the outcomes of evolutionary interactions between plants and their specialized herbivores (Bernays and Graham 1988, Camara 1997, Petschenka and Agrawal 2016).

In conclusion, we have demonstrated that monarch butterflies show substantial genetic variation within and between populations for cardenolide sequestration. The evolution of toxin sequestration in monarchs and other taxa is likely shaped by both evolutionary history (including shifting dietary associations) and contemporary species interactions. Our research also highlights the utility of “natural experiments”—both the monarch’s recent global range expansion and the recent extirpation of birds from Guam— for testing fundamental hypotheses in ecology and evolution.

## Data Accessibility Statement

All raw data and code used in analysis are available through Github at this link: https://github.com/micahfreedman/manuscripts/tree/master/Cardenolide_Sequestration. Data and code are also available through Dryad at this link: https://datadryad.org/stash/dataset/doi:10.25338/B8TD1F.

