## Supplemental Tables S1-S12 and Supplemental Figure S1-S13 for "Population-specific patterns of toxin sequestration in monarch butterflies from around the world"

| Species | Pop | Location | Year | Plant n | Plant Mean | Butterfly n | Butterfly Mean |
|---|---|---|---|---|---|---|---|
| *G. physocarpus* (Hawaii) | 1 | Maui, Hawaii | 2016 | 1 | 4.35 | 8 | 4.58 ± 0.34 |
| | 2 | Maui, Hawaii | 2016 | 4 | 8.27 ± 2.19 | 10 | 6.94 ± 1.10 |
| | 3 | Maui, Hawaii | 2016 | 4 | 8.07 ± 2.85 | 6 | 5.33 ± 1.17 |
| | 4 | Maui, Hawaii | 2016 | 2 | 6.55 ± 0.31 | 6 | 3.73 ± 1.00 |
| | 5 | Maui, Hawaii | 2016 | 3 | 6.80 ± 1.93 | 12 | 5.49 ± 0.27 |
| | 6 | Maui, Hawaii | 2016 | 2 | 14.94 ± 1.00 | 6 | 6.62 ± 0.98 |
| | 10 | Maui, Hawaii | 2017 | 2 | 9.65 ± 2.47 | 8 | 5.80 ± 0.35 |
| | 11 | Maui, Hawaii | 2017 | 3 | 14.38 ± 2.86 | 8 | 6.20 ± 0.73 |
| *G. physocarpus* (Australia) | C1 | Canungra, QLD | 2016 | 2 | 6.78 ± 2.48 | 7 | 4.55 ± 0.50 |
| | C2 | Canungra, QLD | 2016 | 5 | 9.97 ± 1.33 | 11 | 5.20 ± 0.42 |
| | D | Wolffdene, QLD | 2016 | 2 | 9.33 ± 5.94 | 9 | 4.63 ± 0.35 |
| | S | Samford, QLD | 2016 | 4 | 7.44 ± 2.36 | 7 | 4.74 ± 0/70 |
| | N1 | Wisemans Ferry, NSW | 2016 | 1 | 5.37 | 2 | 4.92 ± 1.25 |
| | N2 | Wisemans Ferry, NSW | 2016 | 3 | 7.64 ± 0.57 | 11 | 5.91 ± 0.65 |
| *A. curassavica* (Mariana Islands) | G | Dededo, Guam | 2015 | 19 | 8.87 ± 0.98 | 51 | 11.62 ± 0.60 |
| | GS | Dededo, Guam | 2015 | 4 | 7.23 ± 1.25 | 13 | 11.34 ± 1.15 |
| | R | Songsong, Rota | 2015 | 8 | 9.42 ± 1.25 | 19 | 14.20 ± 1.42 |
| *A. incarnata* (Eastern North America) | 1 | Washtenaw Co., MI | 2016 | 4 | 0.06 ± 0.05 | 12 | 0.50 ± 0.11 |
| | 2 | Washtenaw Co., MI | 2016 | 1 | 0.12 | 4 | 0.53 ± 0.14 |
| | 3 | Washtenaw Co., MI | 2016 | 7 | 0.06 ± 0.03 | 13 | 0.43 ± 0.10 |
| | 4 | Washtenaw Co., MI | 2016 | 2 | 0.22 ± 0.21 | 8 | 0.34 ± 0.14 |
| | 5 | Washtenaw Co., MI | 2016 | 6 | 0.19 ± 0.09 | 17 | 0.34 ± 0.07 |
| *A. syriaca* (Eastern North America) | C1 | Washtenaw Co., MI | 2016 | 3 | 0.55 ± 0.12 | 5 | 2.10 ± 0.78 |
| | C13 | Washtenaw Co., MI | 2016 | 3 | 0.18 ± 0.09 | 7 | 5.37 ± 1.30 |
| | C15 | Washtenaw Co., MI | 2016 | 3 | 0.31 ± 0.05 | 7 | 3.37 ± 0.96 |
| | C21 | Washtenaw Co., MI | 2016 | 3 | 0.57 ± 0.21 | 10 | 6.98 ± 1.81 |
| | C39 | Washtenaw Co., MI | 2016 | 2 | 0.36 ± 0.15 | 9 | 4.49 ± 1.26 |
| | C46 | Washtenaw Co., MI | 2016 | 1 | 0.21 | 4 | 5.18 ± 1.36 |
| | ITH1 | Ithaca, NY | 2012 | 7 | 0.49 ± 0.11 | 21 | 7.27 ± 0.73 |
| | ITH2 | Ithaca, NY | 2017 | 6 | 0.31 ± 0.07 | 11 | 8.98 ± 1.36 |
| | CA | Yolo Co., CA | 2018 | 2 | 0.66 ± 0.50 | 5 | 4.03 ± 1.51 |

| | | | | | | | |
|---|---|---|---|---|---|---|---|
| *A. speciosa* (Western North America) | SN | Seed Needs LLC | 2018 | 16 | 0.21 ± 0.04 | 53 | 3.22 ± 0.43 |
| *A. fascicularis* (Western North America) | Y | Yolo Co., CA | 2012 | 18 | 0.16 ± 0.04 | 57 | 0.31 ± 0.03 |

**Table S1** – Summary of host plant species used in reciprocal rearing experiment. Location refers to the provenance of host plants from which seed was collected. Plant and butterfly sample sizes refer only to the number of individuals from which cardenolides were measured; a larger number of plants were grown and used for measuring larval growth rate (see Freedman et al. 2020, *Evolution*). Plant and butterfly means reflect total cardenolide concentrations (mg/g dry weight) plus or minus mean standard errors.

| Population | Family ID | GOPH | ASCU | AINC | ASFA | ASYR | ASPEC | Total |
|---|---|---|---|---|---|---|---|---|
| **Australia** | AU_1_2017 | 9 | 3 | 0 | 2 | 3 | 0 | 17 |
| | AU_A1_2018 | 3 | 0 | 1 | 1 | 1 | 2 | 8 |
| | AU_A10_2018 | 1 | 0 | 1 | 0 | 1 | 0 | 3 |
| | AU_A11_2018 | 1 | 0 | 0 | 0 | 0 | 0 | 1 |
| | AU_A12_2018 | 2 | 2 | 1 | 1 | 2 | 2 | 10 |
| | AU_A13_2018 | 1 | 2 | 1 | 0 | 1 | 2 | 7 |
| | AU_A14_2018 | 1 | 0 | 1 | 1 | 0 | 1 | 4 |
| | AU_A2_2018 | 1 | 2 | 2 | 2 | 1 | 1 | 9 |
| | AU_A5_2018 | 2 | 1 | 1 | 0 | 2 | 0 | 6 |
| | AU_A8_2018 | 1 | 0 | 1 | 0 | 1 | 0 | 3 |
| | AU_A9_2018 | 1 | 1 | 0 | 1 | 1 | 0 | 4 |
| | | **23** | **11** | **9** | **8** | **13** | **8** | **72** |
| **Population** | **Family ID** | **GOPH** | **ASCU** | **AINC** | **ASFA** | **ASYR** | **ASPEC** | **Total** |
| **California** | CA_1017.1B_2018 | 0 | 1 | 0 | 0 | 1 | 0 | 2 |
| | CA_11_2018 | 1 | 1 | 0 | 0 | 0 | 1 | 3 |
| | CA_13_2017 | 1 | 0 | 0 | 1 | 0 | 0 | 2 |
| | CA_13_2018 | 0 | 1 | 1 | 2 | 1 | 2 | 7 |
| | CA_2_2017 | 1 | 1 | 0 | 2 | 1 | 0 | 5 |
| | CA_2_2018 | 2 | 1 | 1 | 1 | 1 | 2 | 8 |
| | CA_23_2018 | 0 | 1 | 1 | 1 | 1 | 1 | 5 |
| | CA_3_2018 | 5 | 2 | 2 | 2 | 1 | 3 | 15 |
| | CA_5_2017 | 1 | 2 | 0 | 1 | 1 | 0 | 5 |
| | CA_605.1_2018 | 0 | 1 | 1 | 0 | 1 | 1 | 4 |
| | CA_7_2017 | 1 | 0 | 0 | 0 | 1 | 0 | 2 |
| | CA_7_2018 | 1 | 1 | 0 | 1 | 2 | 1 | 6 |
| | CA_8_2017 | 1 | 1 | 0 | 0 | 2 | 0 | 4 |
| | CA_9_2017 | 1 | 0 | 0 | 1 | 1 | 0 | 3 |
| | CA_946.1_2018 | 0 | 0 | 0 | 1 | 0 | 0 | 1 |
| | CA_X3_2018 | 3 | 1 | 2 | 2 | 1 | 2 | 11 |
| | | **18** | **14** | **8** | **15** | **15** | **13** | **83** |
| **Population** | **Family ID** | **GOPH** | **ASCU** | **AINC** | **ASFA** | **ASYR** | **ASPEC** | **Total** |
| **Eastern North America** | ENA_1_2017 | 4 | 3 | 0 | 1 | 2 | 0 | 10 |
| | ENA_2_2017 | 2 | 2 | 0 | 2 | 0 | 0 | 6 |
| | ENA_47_2017 | 1 | 0 | 0 | 1 | 0 | 0 | 2 |
| | ENA_56_2017 | 1 | 1 | 0 | 1 | 2 | 0 | 5 |
| | ENA_70_2017 | 1 | 1 | 0 | 0 | 0 | 0 | 2 |
| | ENA_A_2018 | 1 | 2 | 1 | 1 | 1 | 2 | 8 |
| | ENA_B_2018 | 2 | 1 | 2 | 1 | 3 | 1 | 10 |
| | ENA_C_2018 | 1 | 1 | 2 | 1 | 0 | 1 | 6 |
| | ENA_P_2018 | 2 | 2 | 2 | 1 | 3 | 3 | 13 |
| | ENA_Q_2018 | 3 | 1 | 2 | 1 | 2 | 2 | 11 |
| | ENA_S_2018 | 2 | 1 | 2 | 1 | 0 | 1 | 7 |
| | | **20** | **15** | **11** | **9** | **15** | **10** | **80** |
| **Population** | **Family ID** | **GOPH** | **ASCU** | **AINC** | **ASFA** | **ASYR** | **ASPEC** | **Total** |
| Guam | GU_10_2018 | 0 | 1 | 0 | 0 | 0 | 0 | 1 |
| | GU_13_2018 | 0 | 1 | 0 | 1 | 0 | 1 | 3 |
| | GU_19_2018 | 2 | 2 | 1 | 1 | 2 | 2 | 10 |
| | GU_21_2018 | 1 | 1 | 0 | 0 | 0 | 0 | 2 |
| | GU_25_2018 | 2 | 1 | 1 | 1 | 1 | 1 | 7 |
| | GU_26_2018 | 2 | 1 | 1 | 1 | 1 | 0 | 6 |
| | GU_30_2018 | 0 | 1 | 0 | 0 | 1 | 0 | 2 |

| | | | | | | | | |
|---|---|---|---|---|---|---|---|---|
| | GU_32_2018 | 1 | 1 | 0 | 0 | 1 | 0 | 3 |
| | GU_38_2018 | 2 | 1 | 1 | 1 | 1 | 2 | 8 |
| | GU_40_2018 | 1 | 1 | 1 | 0 | 0 | 1 | 4 |
| | GU_41_2018 | 1 | 1 | 1 | 0 | 1 | 1 | 5 |
| | GU_43_2018 | 3 | 2 | 2 | 1 | 2 | 2 | 12 |
| | GU_901.1B_2018 | 1 | 1 | 1 | 0 | 0 | 0 | 3 |
| | | **16** | **15** | **9** | **6** | **10** | **10** | **66** |
| **Population** | **Family ID** | **GOPH** | **ASCU** | **AINC** | **ASFA** | **ASYR** | **ASPEC** | **Total** |
| Hawaii | HI_1_2018 | 3 | 2 | 2 | 2 | 2 | 2 | 13 |
| | HI_10_2017 | 2 | 3 | 0 | 1 | 3 | 0 | 9 |
| | H1_17_2018 | 0 | 1 | 0 | 1 | 1 | 0 | 3 |
| | Hi_19_2017 | 0 | 1 | 0 | 1 | 1 | 0 | 3 |
| | HI_19_2018 | 2 | 1 | 1 | 1 | 0 | 1 | 6 |
| | HI_2_2018 | 3 | 2 | 2 | 1 | 1 | 3 | 12 |
| | HI_20_2017 | 1 | 0 | 0 | 1 | 0 | 0 | 2 |
| | HI_21_2018 | 2 | 0 | 1 | 1 | 0 | 0 | 4 |
| | HI_22_2017 | 6 | 3 | 0 | 1 | 2 | 0 | 12 |
| | HI_5_2017 | 1 | 1 | 0 | 1 | 0 | 0 | 3 |
| | HI_5_2018 | 0 | 0 | 1 | 0 | 0 | 0 | 1 |
| | HI_944.2_2018 | 1 | 0 | 2 | 0 | 0 | 1 | 4 |
| | | **21** | **14** | **9** | **11** | **10** | **7** | **72** |
| **Population** | **Family ID** | **GOPH** | **ASCU** | **AINC** | **ASFA** | **ASYR** | **ASPEC** | **Total** |
| Puerto Rico | PR_103_2018 | 2 | 1 | 1 | 2 | 1 | 1 | 8 |
| | PR_105_2018 | 2 | 2 | 0 | 0 | 2 | 1 | 7 |
| | PR_107_2018 | 1 | 2 | 2 | 0 | 2 | 2 | 9 |
| | PR_109_2018 | 2 | 0 | 0 | 0 | 1 | 1 | 4 |
| | PR_111_2018 | 2 | 2 | 2 | 2 | 2 | 2 | 12 |
| | PR_112_2018 | 2 | 2 | 1 | 2 | 0 | 1 | 8 |
| | PR_113_2018 | 0 | 1 | 2 | 0 | 1 | 0 | 4 |
| | PR_P1_2018 | 0 | 1 | 0 | 0 | 0 | 0 | 1 |
| | PR_PM2_2018 | 1 | 1 | 0 | 0 | 0 | 0 | 2 |
| | PR_PM4_2018 | 2 | 2 | 2 | 2 | 2 | 2 | 12 |
| | | **14** | **14** | **10** | **8** | **11** | **10** | **67** |

**Table S2 –** Number of wing cardenolide samples for each maternal family, separated by milkweed species. Families are arranged by source population. Note that for monarchs reared in 2017, only four milkweed species (GOPH, ASCU, ASYR, ASFA) were available. In total, we analyzed cardenolides from 440 individual monarchs. Milkweed species abbreviations are as follows: GOPH = *Gomphocarpus physocarpus*, ASCU = *Asclepias curassavica*, AINC = *Asclepias incarnata*, ASFA = *Asclepias fascicularis*, ASYR = *Asclepias syriaca*, ASPEC = *Asclepias speciosa*.

| Compound | Retention Time (Minutes) | Species | Absorbance Peak (nm) |
|---|---|---|---|
| Aspecioside | 1.120 | ASYR, ASPEC | 219.13 |
| Frugoside | 5.933 | ASCU, GOPH, AINC, ASFA | 220.12 |
| Calotropin | 6.660 | ASCU, GOPH | 218.99 |
| Calactin | 7.443 | ASCU, GOPH | 218.77 |
| Digitoxin (Internal Standard) | 10.693 | All samples | 219.20 |

**Table S3** – Cardenolides present in the current study whose identities could be verified with authentic standards. Frugoside was only recorded from AINC and ASFA in trace amounts and may reflect small amounts sequestered by neonate larvae during their first ~12 hours of development on ASCU cuttings, prior to being transferred onto their focal host plants. All compounds provided by A. Agrawal and C. Duplais with the exception of digitoxin (Sigma-Aldrich).

| Wing Wear Score | Criteria Used for Classification | Guam (N) | Rota (N) |
|---|---|---|---|
| **1** | pristine, likely emerged within 1-2 days, no noticeable signs of wing wear | 14 (25.9%) | 5 (18.5%) |
| **2** | margins intact (<=1 pieces missing from margin), some loss of scales on wings but only in small areas | 24 (44.4%) | 12 (44.4%) |
| **3** | chunks (<=4) missing from wing margins; noticeable scale loss leading to reduction in color saturation | 8 (14.8%) | 3 (11.1%) |
| **4** | many chunks (=>5) missing from wing margins, looks fairly battered and has substantial scale loss | 7 (13.0%) | 4 (14.8%) |
| **5** | large chunks missing, extensive scale loss, flight ability likely compromised | 1 (1.8%) | 1 (3.7%) |

**Table S4 –** Criteria used for classifying wild-caught monarchs from Guam and Rota into wing wear categories. Wings were scored based on flatbed scanner images displaying all four monarch wings (both fore- and hindwings), but only hindwings were used for classification, as these were the tissue used for sampling cardenolides. Wing wear values were treated as continuous predictors in models of hindwing cardenolides across islands. Sample sizes within each classification category are shown at right for Guam (n = 54) and Rota (n = 27), as well as the proportion of observations within each category. Mean levels of wing wear were slightly higher on Rota (2.36) than on Guam (2.20).

| Species | Sum of Squares | F | p |
|---|---|---|---|
| GOPH | 13.27 | 103.5 | <0.001 |
| ASCU | 7.72 | 57.2 | <0.001 |
| AINC | 3.93 | 15.1 | <0.001 |
| ASFA | 4.79 | 20.1 | <0.001 |
| ASYR | 7.68 | 27.0 | <0.001 |
| ASPEC | 3.19 | 9.5 | <0.001 |

**Table S5 –** MANOVA results for milkweed species level comparisons of leaf and wing cardenolide profiles. Each row corresponds to a single species-level comparison of leaf and wing cardenolides. Across all species, leaf and wing tissue contained strongly distinct cardenolide profiles, consistent with the idea that sequestration involves active processing of leaf cardenolides.

| Milkweed Species | Compound | Absolute Amount (mg/g) | % of Total Sequestered | PR Amount (mg/g) | Ratio (PR / Others) |
|---|---|---|---|---|---|
| *Asclepias curassavica* | **Frugoside** | 4.190 | 34.4 % | 4.013 | 0.958 |
| | **RT 2.150** | 1.674 | 13.8 % | 4.249 | 2.538 |
| | **RT 0.830** | 1.270 | 10.4 % | 2.015 | 1.587 |
| | **Calotropin** | 1.235 | 10.1 % | 0.689 | 0.558 |
| | **Calactin** | 0.865 | 7.1 % | 0.968 | 1.119 |
| | **RT 5.100** | 0.600 | 4.9 % | 0.598 | 0.997 |
| *Gomphocarpus physocarpus* | **Frugoside** | 1.455 | 26.9 % | 1.887 | 1.297 |
| | **RT 0.830** | 0.785 | 14.5 % | 1.730 | 2.204 |
| | **RT 6.383** | 0.579 | 10.7 % | 0.481 | 0.831 |
| | **Calactin** | 0.564 | 10.4 % | 0.834 | 1.479 |
| | **RT 1.593** | 0.544 | 10.0 % | 1.064 | 1.956 |
| | **Calotropin** | 0.487 | 9.0 % | 0.335 | 0.689 |
| *Asclepias syriaca* | **Aspecioside** | 2.966 | 48.3 % | 0.128 | 0.043 |
| | **RT 1.890** | 0.879 | 14.3 % | 0.098 | 0.111 |
| | **RT 3.590** | 0.744 | 12.1 % | 0.357 | 0.480 |
| | **RT 3.380** | 0.708 | 11.5 % | 0.320 | 0.452 |
| | **RT 2.870** | 0.144 | 2.4 % | 0.031 | 0.215 |
| *Asclepias speciosa* | **Aspecioside** | 1.447 | 44.0 % | 0.096 | 0.066 |
| | **RT 3.380** | 0.592 | 18.0 % | 0.104 | 0.176 |
| | **RT 3.590** | 0.418 | 12.7 % | 0.037 | 0.089 |
| | **RT 1.890** | 0.404 | 12.3 % | 0.363 | 0.899 |
| | **RT 2.870** | 0.070 | 2.2 % | 0.000 | 0.000 |

**Table S6** – Primary sequestered cardenolide peaks across milkweed species, averaged across all monarch populations. The top six compounds are shown for *A. curassavica* and *G. physocarpus*, and the top five compounds are shown for *A. syriaca* and *A. speciosa*. For compounds whose identities are unknown, retention times are listed. Percent of total sequestered refers to within-species totals. In the second column from the right, absolute sequestered amounts are shown for the Puerto Rican population only. The rightmost column shows the ratio of sequestered cardenolides for Puerto Rican monarchs relative to species-level totals across all populations. Note that the ratio for aspecioside sequestered from *A. syriaca* is 0.043, corresponding to 23 times lower sequestration of this compound in Puerto Rican monarchs. For graphical depictions of chromatograms, see Figure 2A. Monarchs reared on *A. incarnata* and *A. fascicularis* contained small amounts of frugoside, RT 6.383, and RT 2.150.

| Predictor | Sum of Squares | $R^2$ | F | DF | p |
|---|---|---|---|---|---|
| Monarch population | 3.42 | 0.033 | 4.77 | 5 | <0.001 |
| Milkweed species | 51.32 | 0.494 | 119.49 | 3 | <0.001 |
| Monarch population x milkweed species | 6.12 | 0.059 | 2.85 | 15 | <0.001 |
| Sex | 0.29 | 0.003 | 2.01 | 1 | 0.066 |
| Residual Error | 42.66 | 0.420 | | | |

**Table S7 –** MANOVA results showing variation explained by milkweed species, monarch population, their interaction, and butterfly sex in the composition of sequestered cardenolides. Compared to quantitative variation in the concentration of sequestered cardenolides (see Table S7), the interaction between monarch population x milkweed species interaction term explained relatively little variation, suggesting that GxE interactions primarily involve variation in the total amount of cardenolide sequestered.

| Model | AIC |
|---|---|
| Wing cardenolides ~ plant cardenolides + species*monarch population + sex + random effects | 760.7 |
| Wing cardenolides ~ species*monarch population + sex + random effects | 756.4 |

**Table S8 –** Comparison between two statistical models of sequestered cardenolides, one that contains individual plant-level cardenolide measurements (top) and one that does not (bottom). The model without individual plant-level measurements of cardenolides has a lower AIC score and is preferred. This, along with data showing an overall weak relationship between plant and wing cardenolide levels within species (Figure S5), suggests that levels of sequestered cardenolides were not strongly determined by available plant cardenolides, at least within milkweed species. The lack of correspondence between leaf and wing cardenolide concentrations may reflect the fact that we deliberately chose to minimize variability in origins of milkweed host plants.

| **Predictor** | **$\chi^2$** | **DF** | **p** |
|---|---|---|---|
| Monarch population | 6.91 | 5 | 0.227 |
| Milkweed species | 61.55 | 3 | <0.001 |
| Monarch population x milkweed species | 77.56 | 15 | <0.001 |
| Sex | 2.85 | 1 | 0.094 |

**Table S9 –** ANOVA results for a linear mixed model comparing total sequestered cardenolide concentrations. Here, the primary term of interest is the interaction between monarch population and milkweed species, which reflects GxE interactions for sequestration ability.

| **Model Term** | **$\chi^2$** | **DF** | **p** |
|---|---|---|---|
| Monarch population | 9.44 | 5 | 0.093 |
| Milkweed species | 79.41 | 3 | <0.001 |
| Sympatric / allopatric status | 0.16 | 1 | 0.687 |
| Sex | 1.34 | 1 | 0.247 |

**Table S10 –** ANOVA results for a linear mixed model directly testing for local adaptation in sequestration ability. As with Table S7, the response variable is total sequestered cardenolides in monarch wings. The primary term of interest is the sympatric/allopatric contrast, which describes the magnitude of performance difference between monarchs reared on sympatric versus allopatric host plants.

| Species | Mean Cardenolide Concentration | Standard Deviation | Coefficient of Variation |
|---|---|---|---|
| GOPH | 5.42 | 2.00 | 0.371 |
| ASCU | 12.17 | 4.83 | 0.397 |
| AINC | 0.45 | 0.45 | 1.002 |
| ASFA | 0.31 | 0.24 | 0.759 |
| ASYR | 6.14 | 4.18 | 0.681 |
| ASPEC | 3.29 | 3.12 | 0.949 |

**Table S11 –** Coefficient of variation in cardenolide sequestration across each milkweed species. Note that variation is lowest on GOPH and ASCU.

| Monarch Population | Milkweed Species | Marginal Mean | SE | df | Lower CL | Upper CL | Group |
|---|---|---|---|---|---|---|---|
| AU | GOPH | 5.242 | 0.795 | 115.159 | 3.667 | 6.816 | A |
| CA | GOPH | 4.819 | 0.885 | 188.960 | 3.074 | 6.564 | A |
| ENA | GOPH | 5.352 | 0.785 | 153.641 | 3.802 | 6.902 | A |
| GU | GOPH | 4.721 | 0.886 | 180.993 | 2.973 | 6.469 | A |
| HI | GOPH | 4.991 | 0.794 | 100.655 | 3.416 | 6.566 | A |
| PR | GOPH | 7.692 | 1.008 | 167.648 | 5.702 | 9.682 | A |
| AU | ASCU | 14.611 | 1.151 | 54.918 | 12.304 | 16.917 | A |
| CA | ASCU | 12.574 | 1.089 | 43.170 | 10.378 | 14.770 | AB |
| ENA | ASCU | 13.079 | 1.021 | 42.408 | 11.018 | 15.139 | A |
| **GU** | **ASCU** | **8.886** | **0.941** | **36.731** | **6.978** | **10.793** | **B** |
| HI | ASCU | 11.013 | 1.052 | 39.936 | 8.887 | 13.140 | AB |
| PR | ASCU | 14.963 | 1.026 | 41.599 | 12.892 | 17.033 | A |
| AU | ASYR | 7.164 | 0.957 | 150.575 | 5.274 | 9.054 | A |
| CA | ASYR | 5.250 | 0.905 | 107.173 | 3.456 | 7.045 | A |
| ENA | ASYR | 6.199 | 0.986 | 99.274 | 4.244 | 8.155 | A |
| GU | ASYR | 6.618 | 1.036 | 177.648 | 4.574 | 8.663 | A |
| HI | ASYR | 7.372 | 1.274 | 108.765 | 4.846 | 9.897 | A |
| **PR** | **ASYR** | **1.056** | **1.040** | **153.891** | **-0.999** | **3.111** | **B** |
| AU | ASPEC | 3.017 | 1.388 | 33.903 | 0.196 | 5.837 | A |
| CA | ASPEC | 4.656 | 1.133 | 23.151 | 2.313 | 6.999 | A |
| ENA | ASPEC | 1.910 | 1.348 | 29.558 | -0.844 | 4.665 | A |
| GU | ASPEC | 3.626 | 1.289 | 33.878 | 1.005 | 6.246 | A |
| HI | ASPEC | 4.955 | 1.481 | 63.087 | 1.994 | 7.915 | A |
| PR | ASPEC | 0.896 | 1.282 | 34.380 | -1.709 | 3.500 | A |

**Table S12** – Estimated marginal means for total sequestered cardenolide concentration for each monarch population x milkweed species. Group level differences were considered significantly different if they had non-overlapping 95% confidence intervals. Combinations of primary interest are shown in bold.

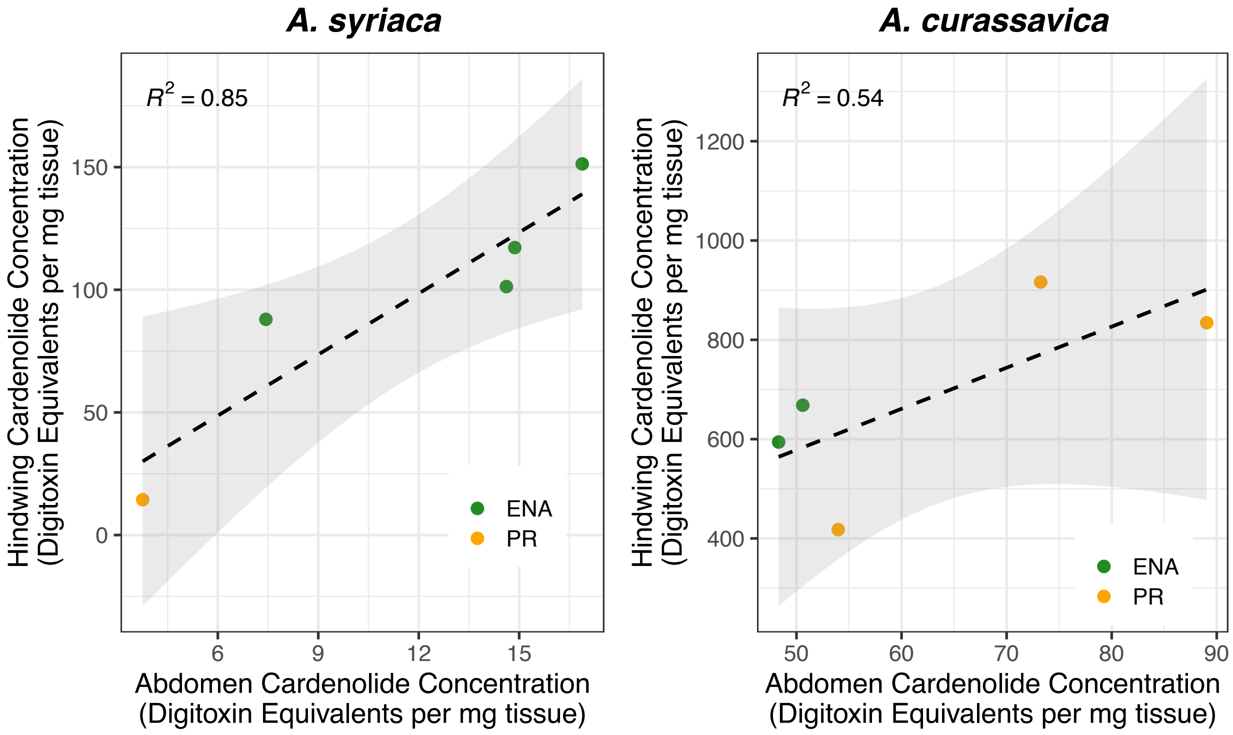


**Figure S1 –** Correlation between cardenolide concentrations in monarch hindwings and abdomens across two host species (*A. syriaca* and *A. curassavica*), with samples from two monarch populations (Eastern North America and Puerto Rico). Hindwing and abdomen cardenolide concentrations are strongly positively correlated within butterflies. Note that correlations shown here correspond to a separate set of butterflies than those described in the main text of the paper. For a full description of methods associated with the butterflies measured in this figure, see Supplementary Appendix 3.

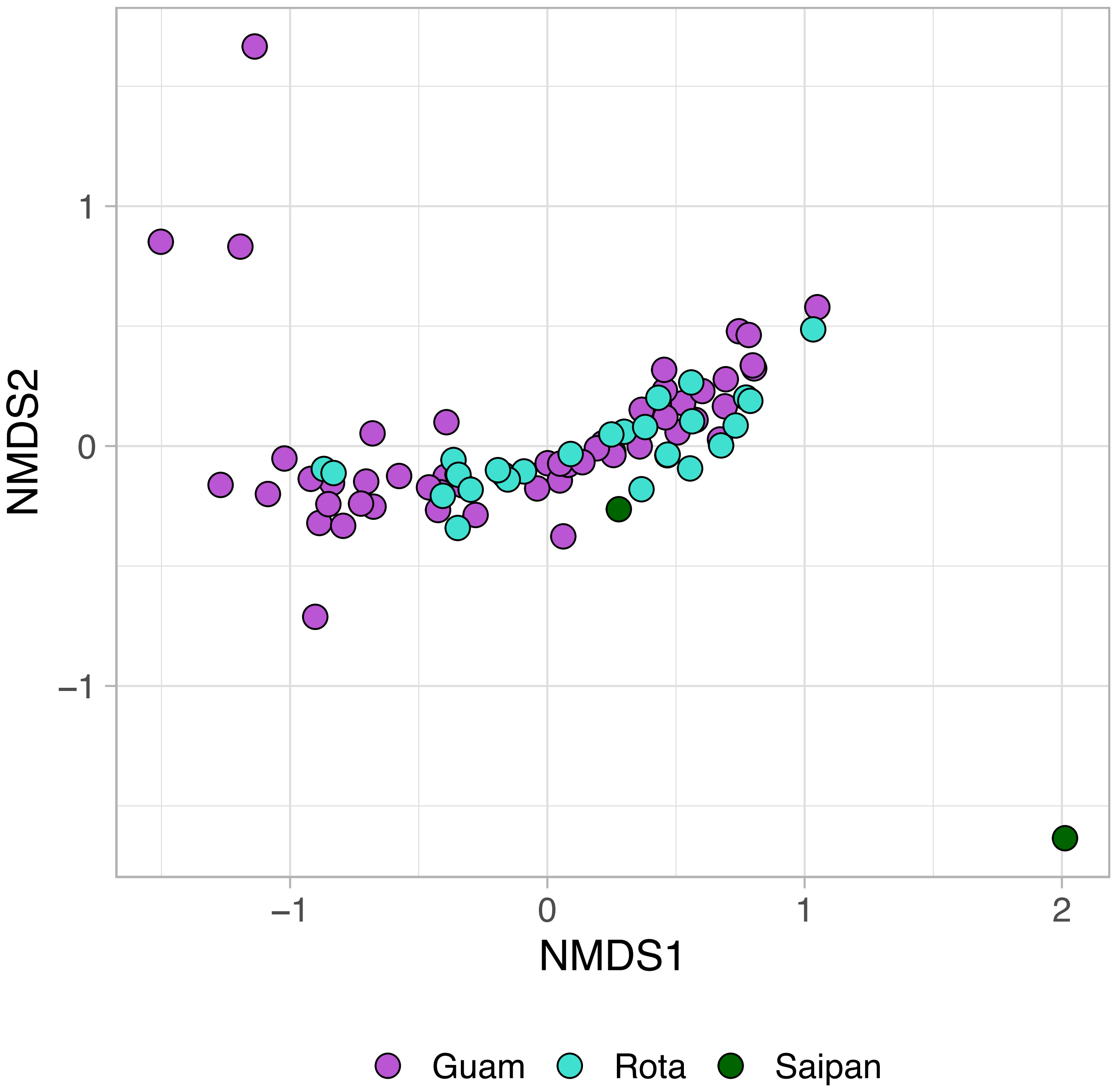


**Figure S2 –** NMDS plot of wing cardenolides from wild-caught monarchs in the Mariana Islands. Butterflies from Guam (n = 54) and Rota (n = 27) generally had indistinguishable cardenolide profiles, consistent with both populations feeding primary on the numerically dominant host *Asclepias curassavica*. Monarchs from Saipan (n = 2) included one wild-caught individual with a cardenolide fingerprint consistent with developing on A. curassavica, as well as one monarch collected on the day of its emergence on an ornamental *Calotropis gigantea* plant (point in lower right).

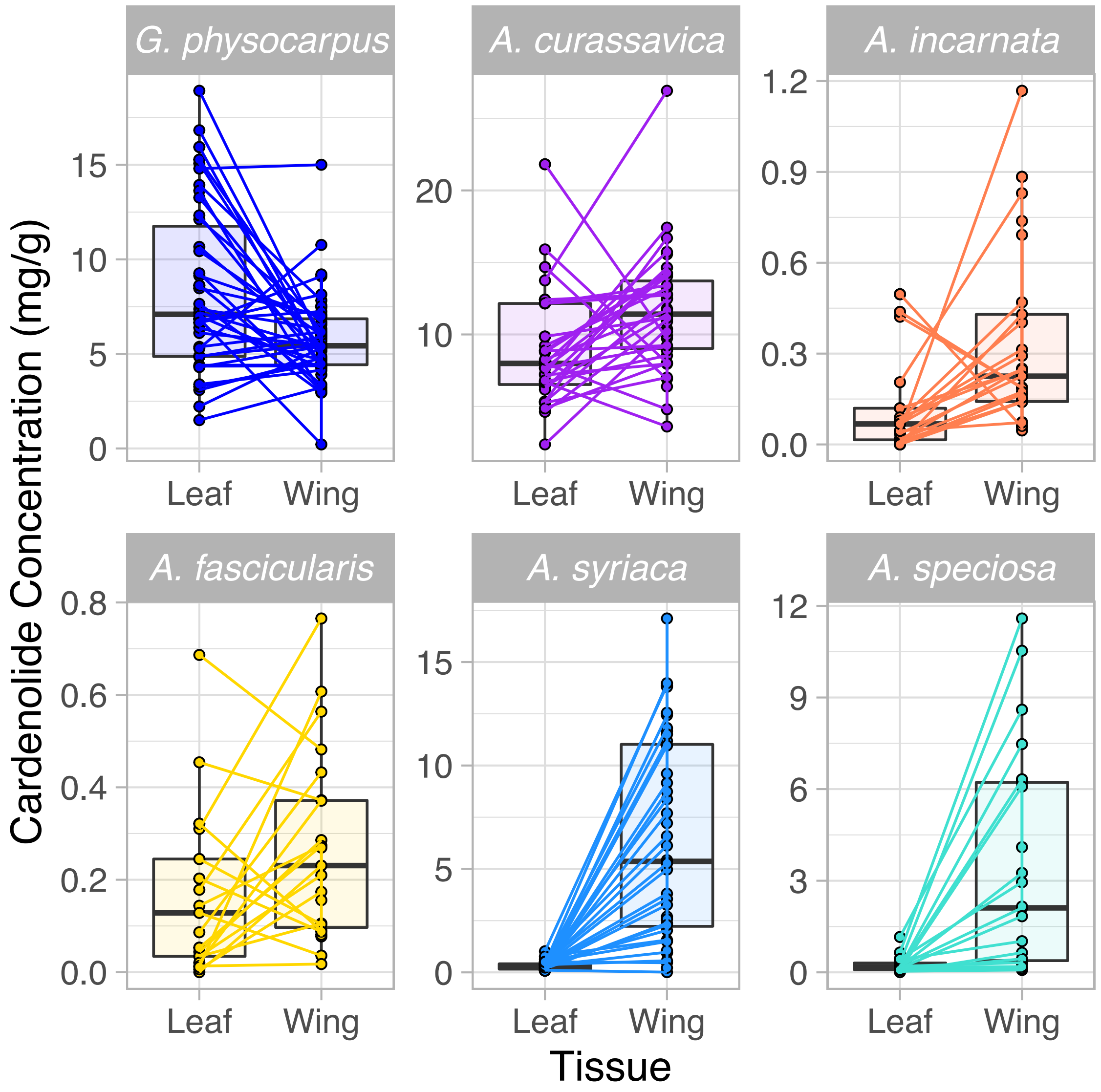


**Figure S3 –** Boxplots showing cardenolide concentrations of milkweed leaf and butterfly hindwing tissue, expressed in mg/g of dry tissue. Here, lines connect individual butterfly samples with their specific natal host plant.

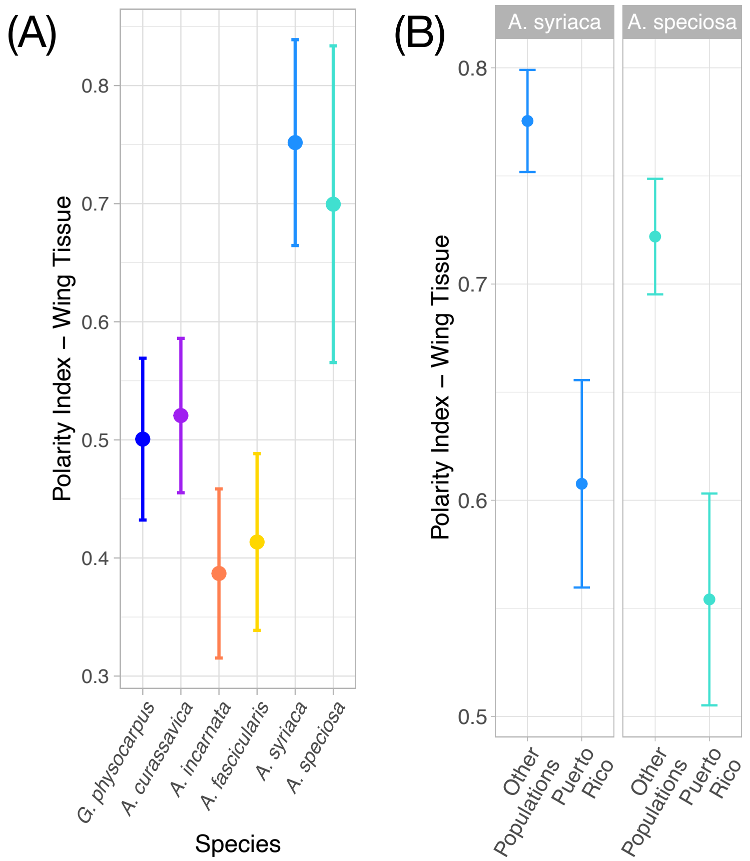


**Figure S4 – (A)** Polarity index of sequestered cardenolides across each milkweed species. Here, a value closer to 1 corresponds to a sequestration profile biased towards compounds with early retention times and high polarity. Compounds sequestered from A. syriaca and A. speciosa were disproportionately polar. **(B)** Compared to all other populations, the subset of sequestered compounds from A. syriaca and A. speciosa was overall less polar. This likely corresponds to the disproportionately low levels of aspecioside (the earliest eluting compound) in hindwings of Puerto Rican monarchs (see Table S5).

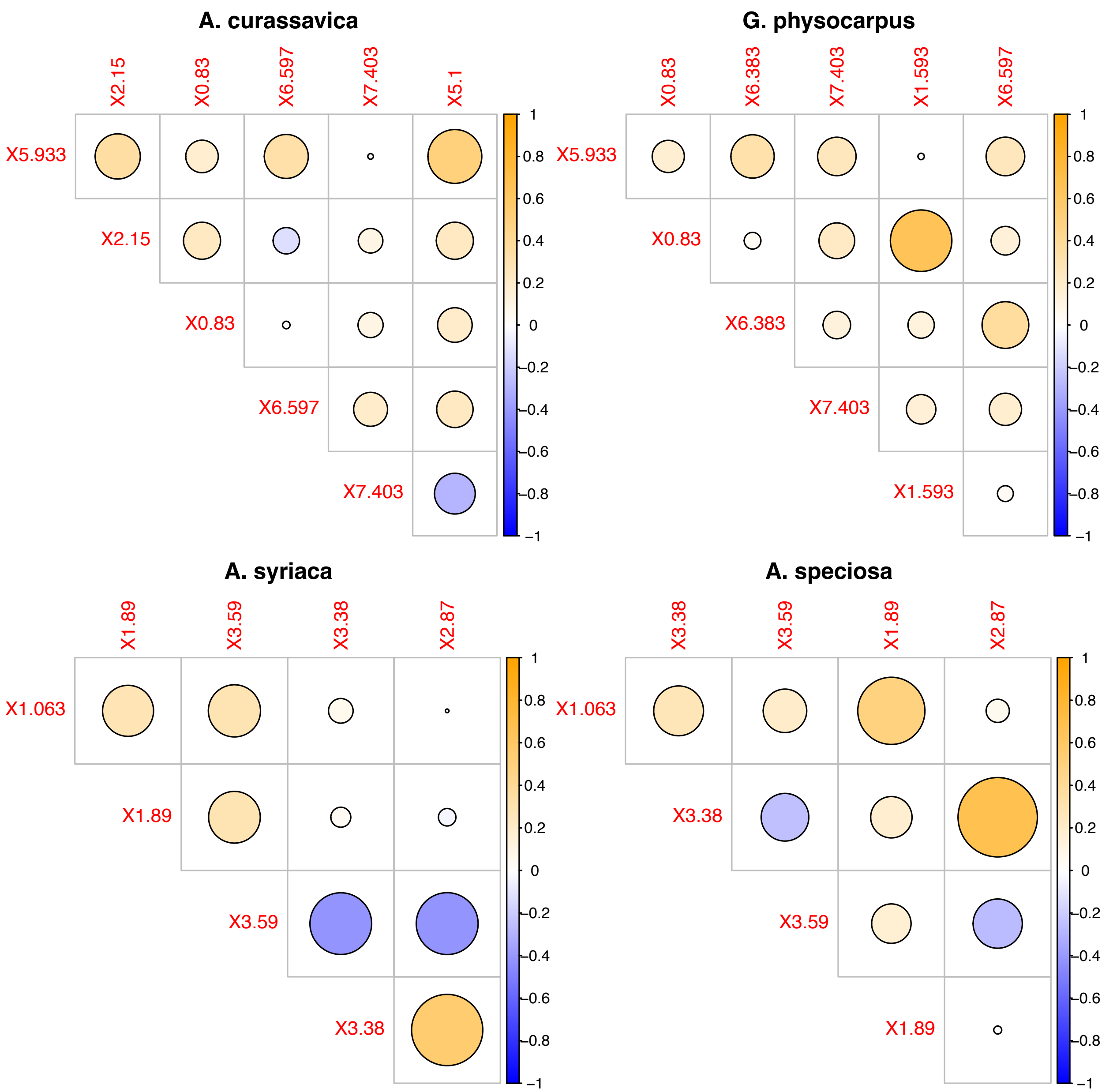


**Figure S5 –** Pairwise correlations among the major compounds sequestered for each of the primary milkweed species of interest. The size of the circle corresponds to the strength of the correlation, with orange dots indicating positive correlations and blue dots indicating negative correlations. Most compounds were positively correlated with each other across butterflies, with a few exceptions. Individual compounds are the same as those reported in Table S5: X5.933 is frugoside, X6.597 is calotropin, X7.403 is calactin, and X1.063 is aspecioside. Numbers in compound names correspond to retention times. Note that only the five most abundant compounds from *A. curassavica* and *G. physocarpus* are shown, and only the four most abundant compounds are shown for *A. syriaca* and *A. speciosa*.

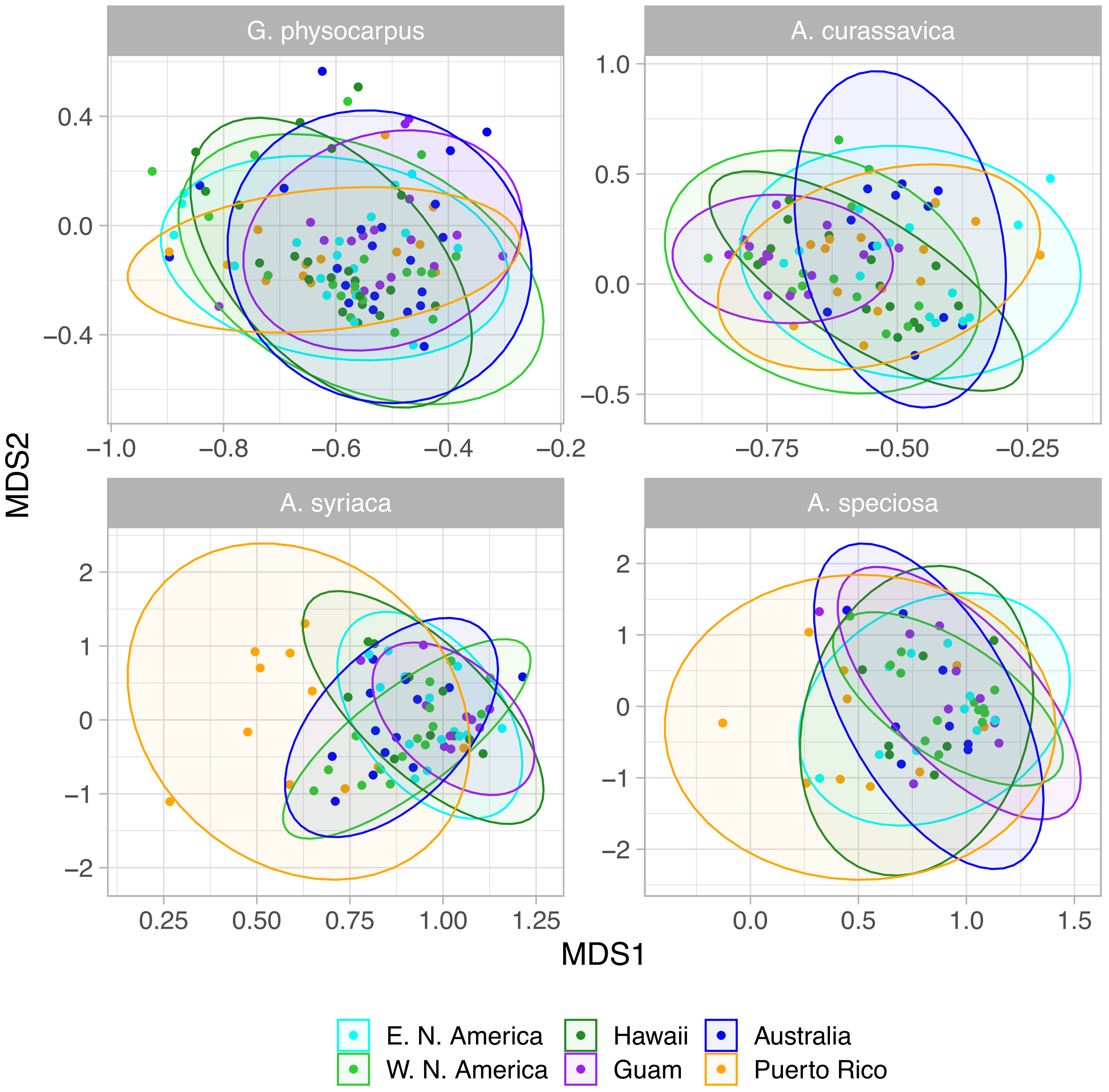


**Figure S6 –** Multivariate disparity in sequestered cardenolide profiles, shown separately for each milkweed species. Results shown are based on a single overall dissimilarity matrix but are faceted by milkweed species. Ellipses correspond to the 95% confidence profiles and were generated using the stat_ellipse function. All populations appear to have generally similar overall multivariate sequestration profiles, with the potential exceptions of Puerto Rican monarchs reared on *A. speciosa* and especially on *A. syriaca*.

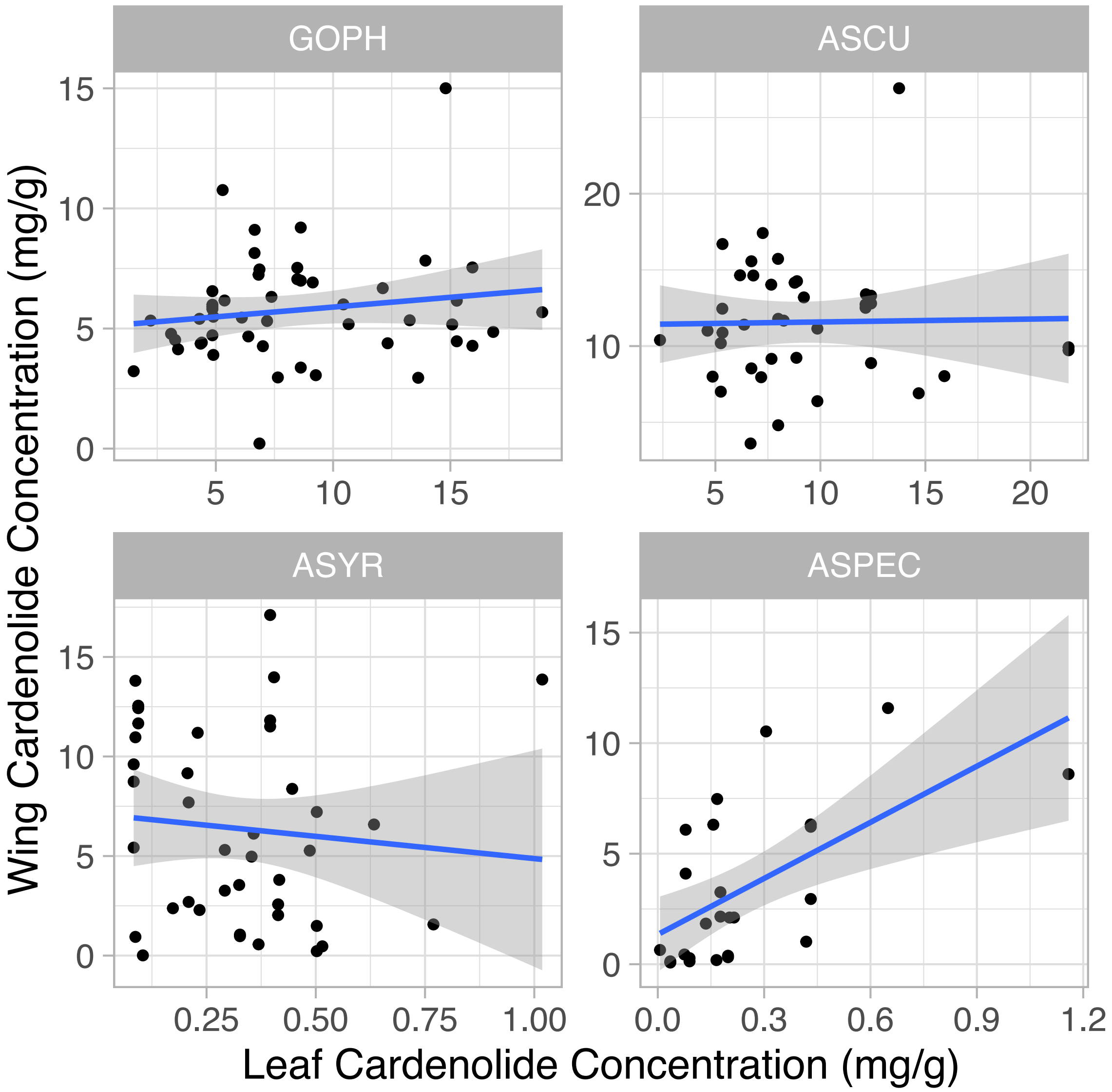


**Figure S7** – Raw data showing correlations between leaf cardenolides and wing cardenolides for the subset of wing samples that had matched plant samples. When comparing butterfly samples whose corresponding natal plant was also analyzed (n = 154), there was little evidence for a correlation between dietary cardenolides and sequestered cardenolides. The lone species with a significantly positive correlation was *A. speciosa* (t = 2.822, p = 0.011).

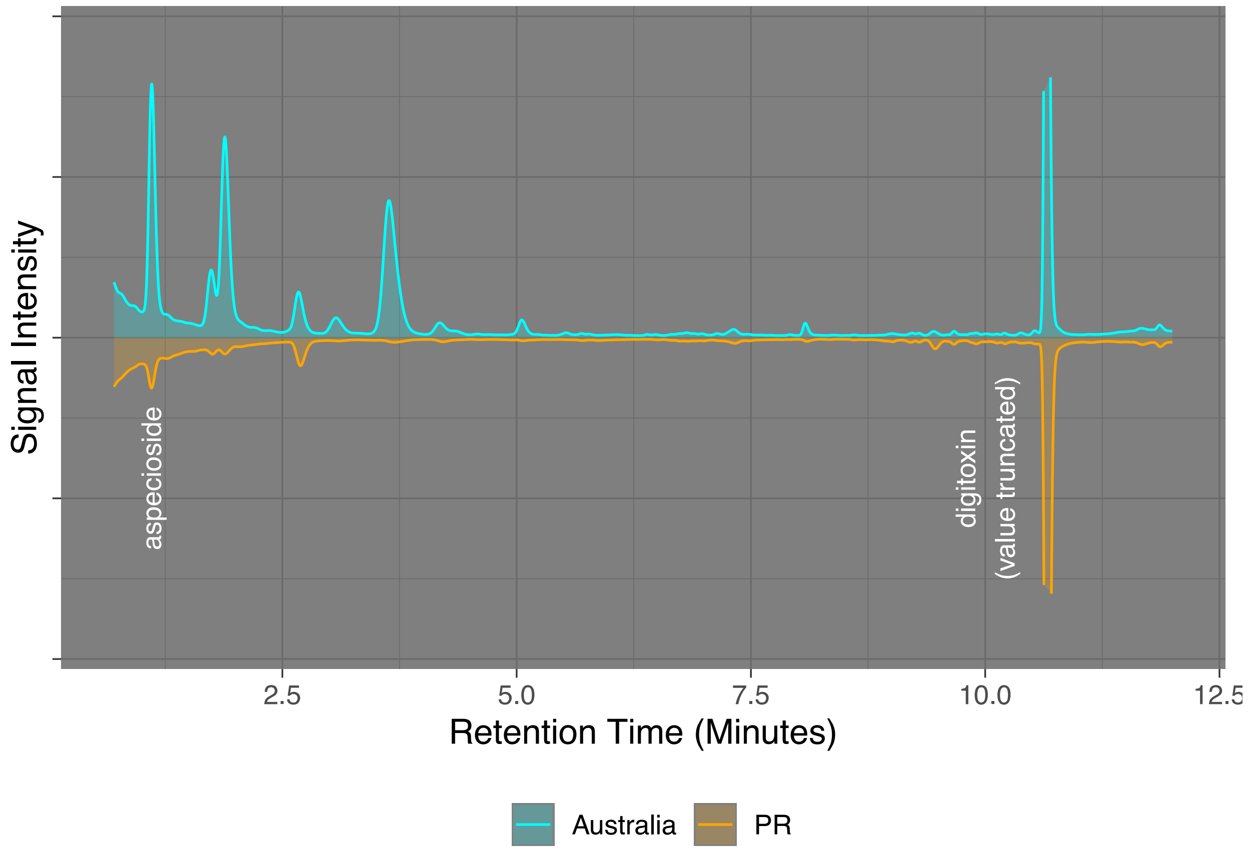


**Figure S8** – Example of chromatograms from two monarchs reared on *A. syriaca*. Australian monarchs (cyan), despite more than 150 years isolated from this ancestral North American host, still retain their ability to sequester normally. By contrast, Puerto Rican monarchs (orange) sequester poorly from *A. syriaca*, with average total cardenolide concentrations that are more than five times lower than all other populations. Sequestration of the compound aspecioside in Puerto Rican populations is especially poor, with average concentrations that are nearly 25 times lower.

(A)

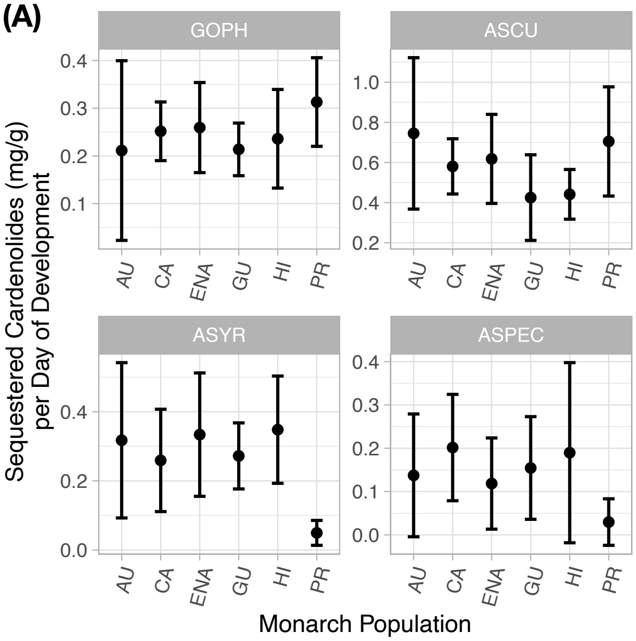


(B)

| Measure | Species | Puerto Rico | Other Populations | Ratio |
|---|---|---|---|---|
| Cardenolide Concentration | GOPH | 7.473 | 5.150 | 1.450 |
| | ASCU | 15.689 | 11.476 | 1.367 |
| | ASYR | 1.203 | 7.036 | 0.171 |
| | ASPEC | 0.768 | 3.810 | 0.201 |
| Development-adjusted Sequestration | GOPH | 0.313 | 0.242 | 1.293 |
| | ASCU | 0.542 | 0.705 | 1.301 |
| | ASYR | 0.050 | 0.306 | 0.162 |
| | ASPEC | 0.030 | 0.162 | 0.183 |

**Figure S9 –** Accounting for development time does not meaningfully impact inferences related to cardenolide sequestration. (A) Cardenolide concentration divided by days from hatching to eclosion, for each *population x species* combination. Note that the results are virtually indistinguishable from those shown in Figure 4A. (B) Comparison of cardenolide concentration and development-adjusted cardenolide sequestration for Puerto Rican monarchs versus all other monarch populations. Puerto Rican monarchs had modestly slower development time than all other populations across all hosts (see Freedman et al. 2020a), but still had the highest levels of sequestration on *A. curassavica* and *G. physocarpus*. Ratio is simply the value for the Puerto Rican population divided by the value for all other populations.

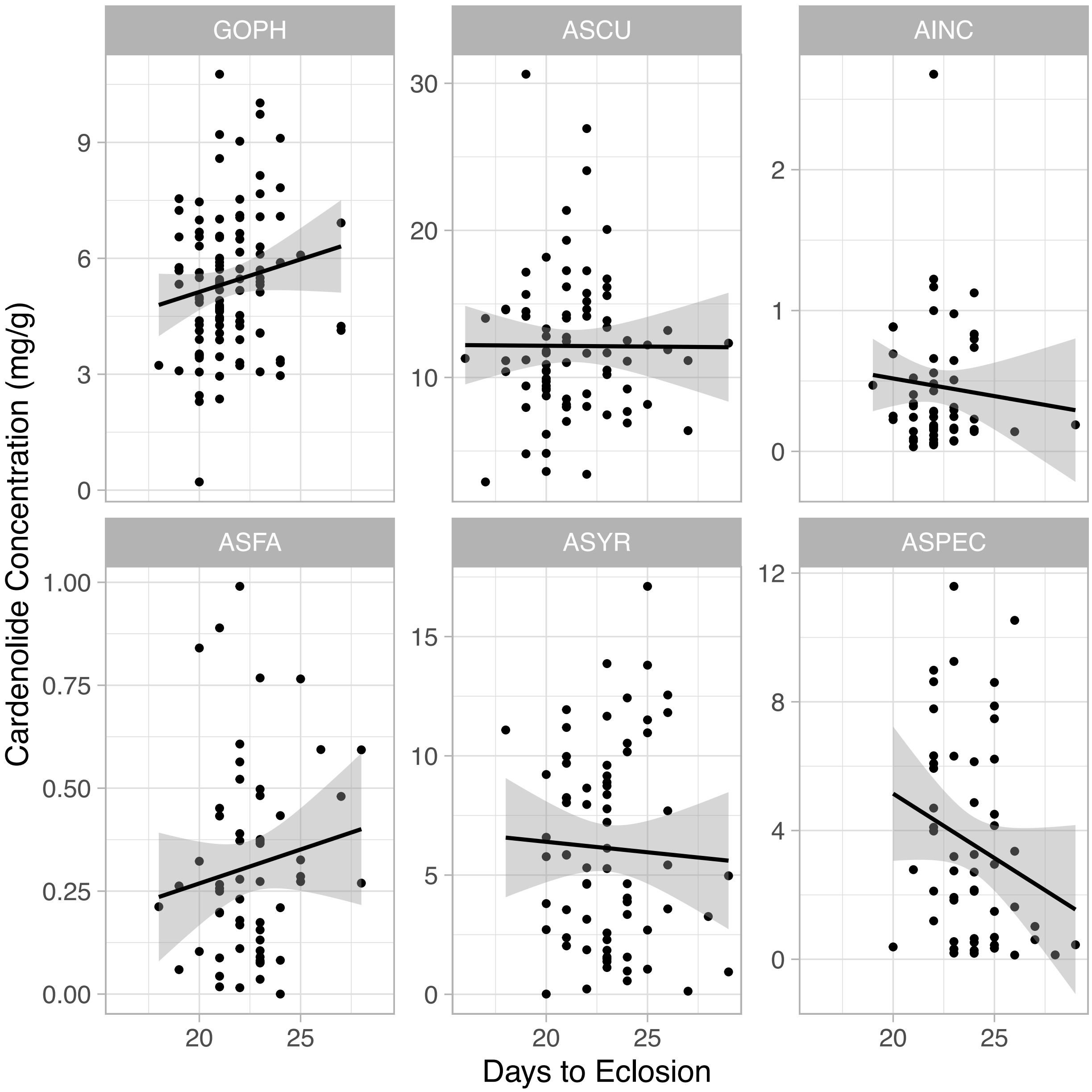


**Figure S10 –** Raw data showing relationship between development rate (measured as days from hatching to eclosion) and cardenolide sequestration. There was no overall correlation between development time (measured as days from egg hatching until eclosion) and the overall quantity of cardenolide sequestered (t = 0.198, p = 0.844).

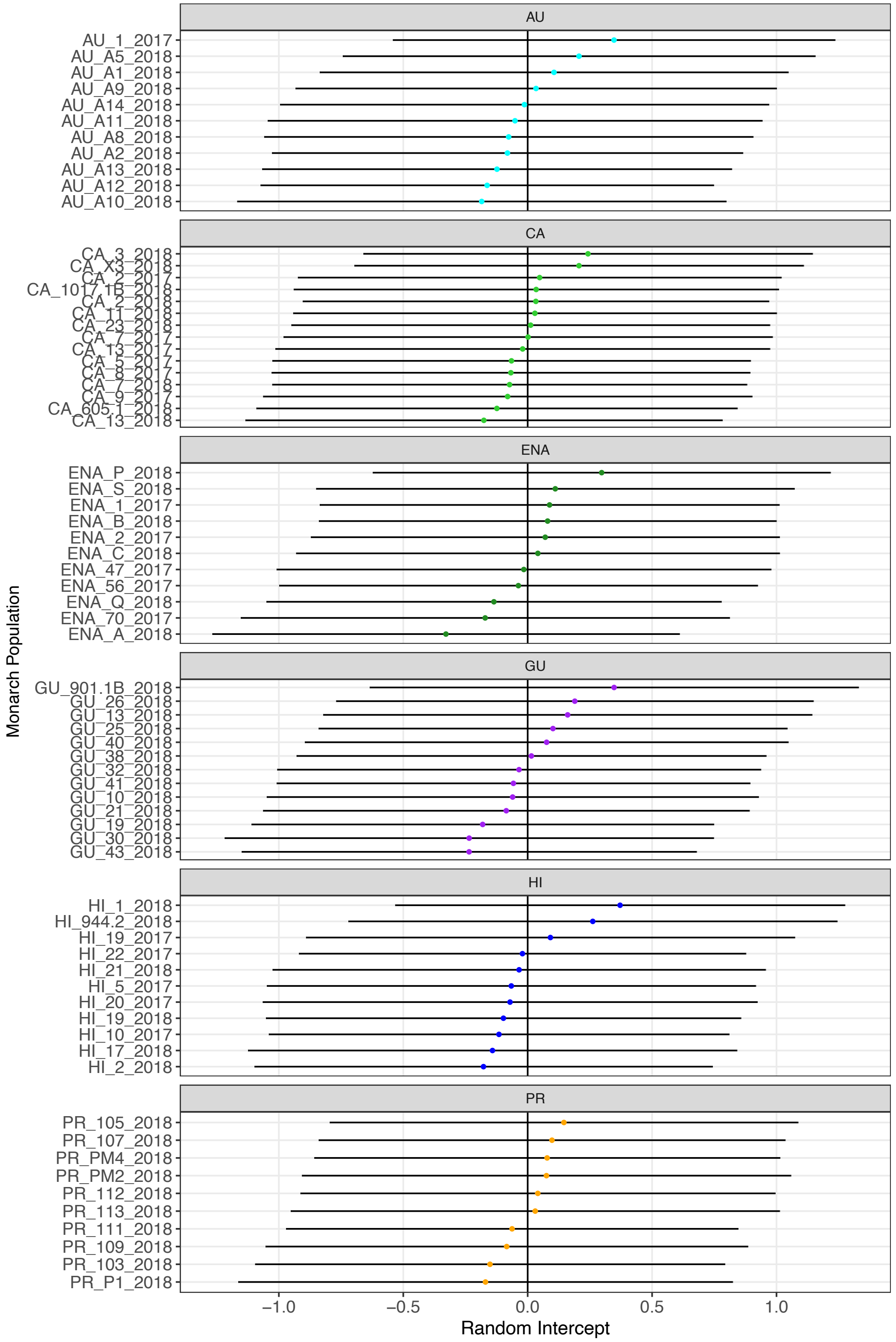


**Figure S11** – Random intercepts and associated 95% confidence intervals from the model testing for GxE interactions in sequestration. Intercepts are shown for each maternal family within each population of interest. Note that intercepts are normally distributed around 0 within each monarch population.

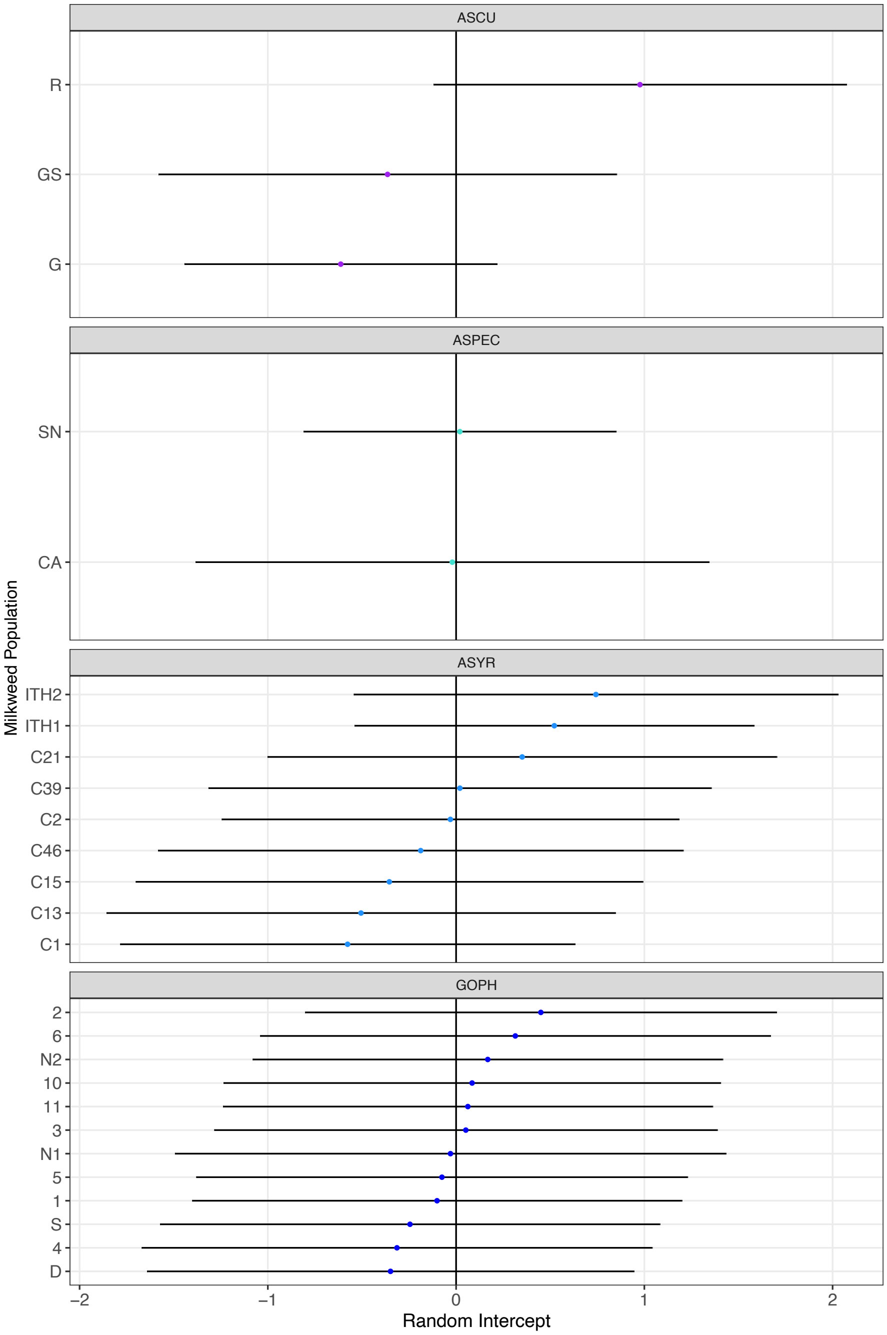


**Figure S12** – Random intercepts and associated 95% confidence intervals from the model testing for GxE interactions in sequestration. Intercepts are shown for each plant population within each milkweed species of interest.

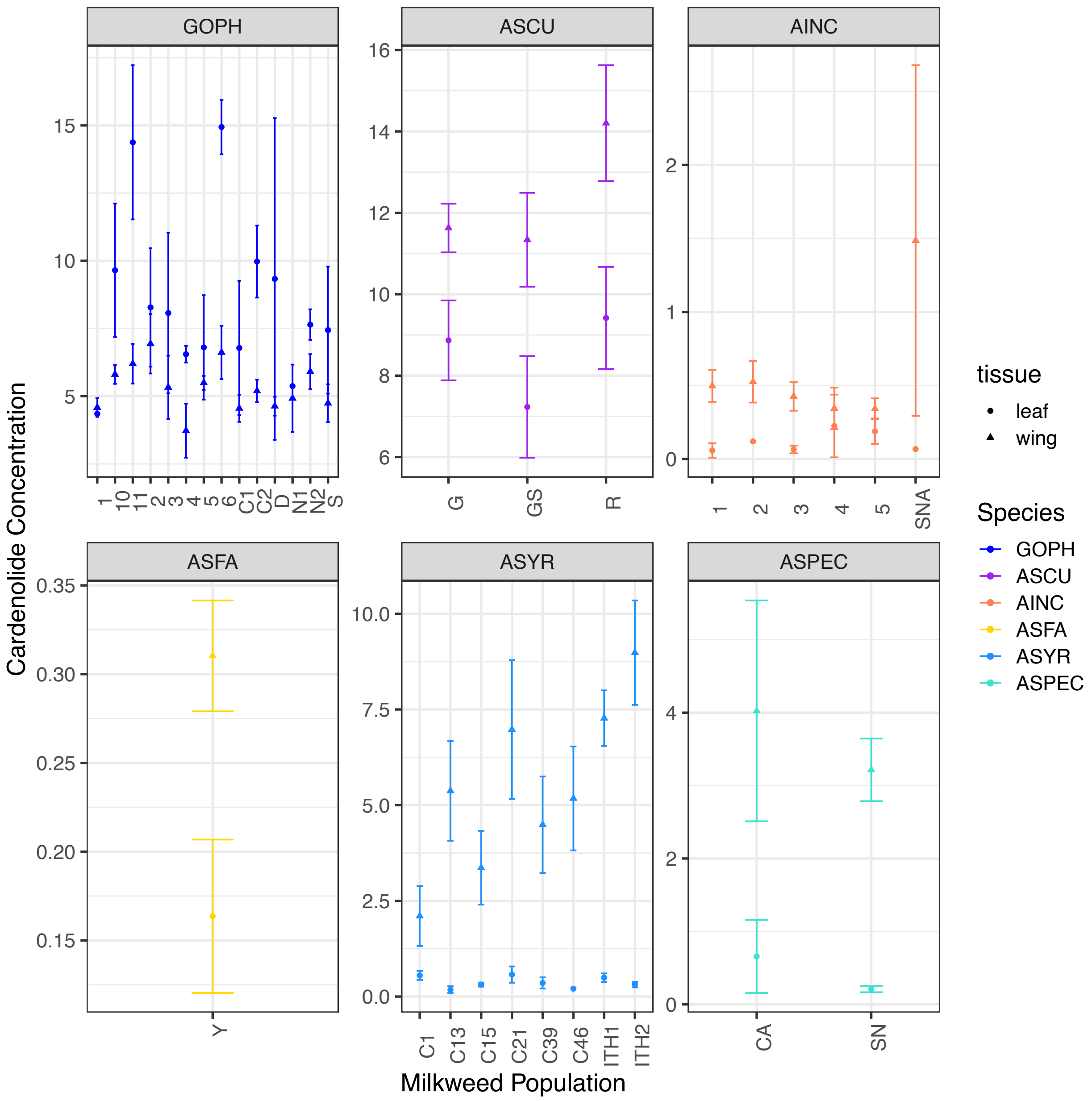


**Figure S13** – Correspondence between sequestered cardenolide (triangles) and leaf tissue (circles) for each plant population within milkweed species. For a disaggregated regression of these relationships, see Figure S7.
